# Intracellular Mechanosensation in Intestinal Smooth Muscle: Piezo1 Complexes Amplify Signaling Beyond the Surface

**DOI:** 10.64898/2026.02.19.706403

**Authors:** Geoanna M. Bautista, Declan Manning, Emily C. Lieu, Collin Matsumoto, Stephanie I. Ugochukwu, Joshua P. Tulman, Miguel Martin-Aragon Baudel, Nuria Daghbouche-Rubio, Steven J. McElroy, Sal A. Baker, Manuel F. Navedo, L. Fernando Santana

## Abstract

Mechanosensation is fundamentally viewed as a plasma membrane phenomenon. We challenge this paradigm by introducing intracellular mechanosensation in intestinal smooth muscle. We hypothesized that a distinct, organelle-based signaling axis exists to amplify mechanotransduction from the inside out. To test this, we investigated whether Piezo1, a canonical plasma membrane mechanosensor, also operates within the cell. Using tissue-level wire myography, high-resolution confocal microscopy, proximity ligation assays, and patch-clamp electrophysiology on freshly dissociated cells, we identified a functional intracellular signaling hub that starts at the sarcoplasmic reticulum (SR). Unlike surface transduction, this intracellular mechanism relies on a nanoscale multiprotein complex (<40 nm) comprising an SR sensor (intra-Piezo1) and an amplifier (Ryanodine Receptor, RyR), coupled with a PM effector (large-conductance, Ca^2+^-activated K^+^ channels, i.e., BK_Ca_ channels). Activating this intracellular complex generated massive BK-mediated outward currents independent of extracellular Ca²⁺ but strictly dependent on internal SR Ca²⁺ stores, confirming intrinsic organellar mechanotransduction. Within this complex, intra-Piezo1 and RyR are positioned to operate as a coupled SR Ca²⁺ release unit that activates BK channels at SR–PM junctions, driving potent membrane hyperpolarization that reduces smooth muscle contractility, revealing the intra-Piezo1 complex as a molecular brake on excitation. These findings demonstrate that mechanotransduction is not confined to the cell surface. Instead, a specialized Sensor–Amplifier–Effector complex originating at intracellular organelles amplifies cellular sensitivity to physical force, providing a critical gain-control system that restrains smooth muscle excitability and regulates GI motility.

**Key Points:**

- Intracellular organelles contribute to mechanosensory signaling in GI smooth muscle cells, complementing plasma membrane mechanisms.
- Intra-Piezo1 form a nanoscale signaling complex (<40nm) on the sarcoplasmic reticulum (SR), linking the mechanosensor Piezo1 with RyR and large conductance, Ca^2+^-activated K^+^ channels.
- Unlike surface sensors, this intracellular complex functions via a “Sensor-Amplifier-Effector” mechanism in which intra-Piezo1 detects mechanical stress and triggers SR Ca^2+^ release, thereby activating a nearby RyR and large-conductance, Ca^2+^-activated K^+^ channel.
- Engaging this intracellular Piezo1-mediated axis significantly dampens smooth muscle contractility, identifying a critical gain-control system essential for regulating GI motility.

## INTRODUCTION

Mechanotransduction is a fundamental physiological process by which cells convert physical forces, such as shear stress, tension, and hydrostatic pressure, into biochemical signals (Huang *et al*., 2004). This conversion is essential for diverse biological functions, ranging from vascular tone regulation to sensory perception. In smooth muscle, mechanical stimuli such as intraluminal pressure or tissue stretch trigger changes in intracellular Ca²⁺ that regulate contractility and gene expression (Hill-Eubanks *et al*., 2011). However, Ca²⁺ signaling in mechanotransduction involves considerable spatiotemporal complexity rather than simple global changes (Earley, 2013).

In the gastrointestinal (GI) tract, mechanosensitivity is essential for coordinating motor patterns that drive peristalsis, regulate luminal flow, and maintain barrier function (Mercado-Perez & Beyder, 2022). The intestinal wall undergoes continuous mechanical activity from luminal distension, peristaltic strain, and fluid shear stress. The intestinal smooth muscles must then integrate these diverse mechanical signals through the SIP syncytium, an electrically coupled network that includes interstitial cells of Cajal and Pdgfrα⁺ cells (Sanders, 2019; Sanders *et al*., 2023), to collectively drive the motor patterns required for intestinal homeostasis.

Central to this process are mechanosensitive ion channels (MSCs), which rapidly gate in response to membrane deformation and convert mechanical forces into electrical signals across diverse physiological systems (Gu & Gu, 2014; Alcaino *et al*., 2017). In GI smooth muscle, these consist of Piezo1 channels and transient receptor potential (TRP) channels, which contribute to depolarizing cation currents, potassium channels like Ca^2+^-activated K^+^ channels (BK_Ca_), and TREK-1 that provide negative feedback to reduce overexcitability (Sanders & Koh, 2006; Gonzalez-Cobos & Trebak, 2010; Berrier *et al*., 2013; Alcaino *et al*., 2017). These channels have been predominantly studied and understood as plasma membrane mechanosensors, positioned to detect extracellular mechanical forces and translate them into electrical and Ca²⁺ signals that regulate contractility via interactions with nearby voltage-gated Na^+^ and Ca^2+^ channels (Sanders, 2008).

This plasma membrane (PM)-centric paradigm underpins current mechanotransduction models in GI smooth muscle. According to this framework, Piezo1 channel activation at the PM would lead to cation influx, membrane depolarization, and the activation of voltage-gated Ca²⁺ channels (Ca_V_1.2) (Joshi *et al*., 2021) This sequence is traditionally viewed as an excitatory cascade: Force → Cation Influx → Contraction, a model supported by data in vascular tissues where Piezo1 activation clearly modulates Ca^2+^-dependent excitation (Lim & Harraz, 2024).

However, Piezo1⁻/⁻ intestinal smooth muscle presents a striking paradox. Rather than simply losing mechanosensitive excitation, these tissues exhibit profound compensatory remodeling: upregulation of Ano1 Ca²⁺-activated Cl⁻ channels and increased Ca_V_1.2 expression drive electrical hyperexcitability, yet contractile force remains significantly reduced compared to wildtype (Bautista et al., 2025). This dissociation of enhanced excitation coupled with diminished contraction cannot be simply explained by the loss of a plasma membrane excitatory channel. If Piezo1 deletion merely removed one source of depolarizing current, compensatory upregulation of other excitatory channels should restore or even enhance contractility. Instead, the tissue becomes more electrically excitable, but contractions become even weaker. This suggests that Piezo1 deletion removes a fundamentally different type of mechanosensory signal, one that is not simply excitatory, implying a more complex role (Bautista *et al*., 2025a; Bautista *et al*., 2025b).

A new body of evidence is challenging the previous paradigm that mechanotransduction is restricted to the cell surface. Early studies identified Piezo1 in the endoplasmic reticulum of epithelial cells (McHugh et al., 2010) and at the endo/sarcoplasmic reticulum (ER/SR)-PM junctions (Zhang et al., 2017). More recently, Liao et al. (2021) provided evidence for intracellular Piezo1 in pulmonary smooth muscle, where it mediates Ca^2+^release from the SR independently of extracellular Ca^2+^. These findings point toward a concept of “intracellular mechanosensation,” in which organelles possess intrinsic capabilities to detect and respond to physical stress. Based on this emerging data and on the functional deficits reported by our group (Bautista *et al*., 2025a), we hypothesized that the primary role of Piezo1 in GI smooth muscle is not to mediate surface influx, but to anchor a specialized intracellular signaling complex. We propose that this complex localizes to the SR and functionally couples the mechanosensor (Piezo1) to intracellular amplification machinery (Ryanodine Receptors and BK channels).

To test this hypothesis, we utilized high-resolution confocal microscopy and proximity ligation assays (PLA) to map the nanoscale architecture of Piezo1 relative to SR calcium release channels in intestinal smooth muscle. We combined these approaches with patch-clamp electrophysiology and wire myography to determine whether activating this complex drives hyperpolarizing currents dependent on internal stores rather than surface entry, thereby altering contractile properties. Our data reveal a novel physiological axis: an “intracellular mechanosensation” pathway in which organelle-based Piezo1 complexes amplify mechanotransduction to fine-tune GI motility.

## MATERIALS AND METHODS

### Animals

All animal procedures were approved by the University of California, Davis Institutional Animal Care and Use Committee (IACUC Protocol #25185) and conducted in accordance with the National Institutes of Health Guide for the Care and Use of Laboratory Animals. Adult male and female C57BL/6J mice (8-12 weeks old, 20-30 g body weight; RRID: IMSR_JAX:000664) were obtained from Jackson Laboratory and housed under standard laboratory conditions (12:12h light-dark cycle, 22±2°C, ad libitum access to standard rodent chow and water). For immunocytochemistry experiments, Piezo1-tdTomato reporter mice (Piezo1tm1.1(KOMP), Jackson Laboratory Stock #029214; RRID: IMSR_JAX:029214) were bred and maintained as described previously (Ranade *et al*., 2014). Mice were euthanized by isoflurane inhalation (5% in O₂) followed by cervical dislocation. Death was confirmed by absence of heartbeat and toe-pinch reflex. Both male and female mice were used, and data were pooled as no sex-specific differences were observed in any measured parameters.

### Tissue Preparation and Cell Dissociation

Following euthanasia, the small intestine was rapidly excised and placed in ice-cold Ca^2+^-free modified Hank’s solution (MHS) containing (in mM): 125 NaCl, 5.36 KCl, 2 MgCl2, 0.336 Na2HPO4, 0.44 K_2_HPO_4_, 11 HEPES, 15.5 NaHCO_3_, 10 Glucose, and 2.9 sucrose. External muscularis layers were peeled and gently separated from the mucosa, then equilibrated in Ca^2+^-free MHS. Freshly dissociated smooth muscle cells were obtained for patch-clamp electrophysiology, immunocytochemistry (ICC), and proximity ligation assays (PLA) as described previously (Koh *et al*., 1998). Briefly, muscle strips were incubated in two successive Ca^2+^-free enzyme solutions containing (per ml): 1) papain (Thermo Fisher, 0.5-1 mg), DTT (Sigma, 1 mg), bovine serum albumin (Sigma, 1 mg) and 2) collagenase (Worthington, Type II, 2 mg), trypsin inhibitor (Worthington, 1 mg), BSA (Sigma, 2 mg) and ATP (Sigma, 0.27 mg).

### Wire Myography

#### Tissue equilibration

Segments of murine small intestine (full-thickness and peeled muscularis, approximately 3 mm in length) were mounted on a wire myograph (DMT820MO, Danish Myo Technology, Denmark) in Krebs-Ringer bicarbonate solution maintained at 37°C and pH 7.40, continuously bubbled with carbogen (95% O2/5% CO2). Following mounting, preparations were allowed to equilibrate for 30 minutes with solution changes every 10 minutes. Preparations that failed to develop spontaneous phasic contractions after equilibration or that showed amplitude <0.05 mN were excluded from analysis. Spontaneous contractile activity was recorded continuously using LabChart software (AD Instruments, Colorado Springs, CO) at a sampling rate of 250 Hz. Force data were exported as tab-delimited text files for offline analysis.

#### Experimental protocols

After establishing stable spontaneous contractions for 5 minutes (steady state baseline), pharmacological agents were added cumulatively to the organ bath. For Piezo1 activation experiments, Yoda2 (Sigma, 25 µM) was applied, and responses were recorded for 30 minutes. Yoda2, a second-generation Piezo1 activator with improved selectivity and potency compared to Yoda1, was chosen for its advantages in *ex vivo* tissue preparations and consistent activation across cell-permeable compartments. For ryanodine receptor experiments, tissue was pretreated with Ryanodine (ABCAM, 100 µM) for 60 minutes prior to Yoda2 application. For SR Ca^2+^ store depletion experiments, tissue was pretreated with Thapsigargin (Fisher, 10 µM) for 15 minutes prior to Yoda2 application. Vehicle controls (0.1% DMSO) showed no significant effects on contractile parameters.

#### Contractile analysis

Contractile force recordings were analyzed using a custom Python-based analysis toolkit (source code available at https://doi.org/10.5281/zenodo.18472624). Raw force traces were imported and processed to detect contractions using automated peak detection (Wire Myography Analyzer v_4; SciPy find_peaks) with validated settings: prominence and height thresholds set at 0.05 mN, a minimum inter-peak distance of 1.0 seconds, and a minimum peak width of 0.3 seconds. Metrics such as frequency (cpm), amplitude (mN), and kinetic parameters (rise/relaxation rates) were derived from standardized 150-s analysis windows. Technical replicates from the same animal were averaged for statistical comparison. Contractile parameters were compared before and after drug application using paired comparisons within the same tissue preparation. Automated validation plots were generated for every recording, displaying the full force trace with marked peaks, the calculated baseline, and the detection threshold. All plots were visually inspected to verify accurate peak detection. Recordings with considerable movement artifacts, unstable baselines, or equipment issues were excluded from analysis.

### Whole-Cell Patch Clamp Electrophysiology

#### Recording configuration

Whole-cell voltage-clamp recordings were obtained from freshly isolated intestinal smooth muscle cells (SMCs) at room temperature using an Axopatch 200B amplifier, a Digidata 1440 digitizer, and Clampex 10 software; data were analyzed with Clampfit (Molecular Devices, Sunnyvale, CA). Borosilicate glass pipettes (P-97 puller, Sutter Instruments, Novato, CA) were polished to a resistance of 2–4 MΩ. Cells were voltage-clamped at a holding potential of -50 mV; currents were sampled at 50 kHz and low-pass filtered at 2 kHz; recordings were acquired in gap-free mode.

#### Solutions

Standard external Tyrode’s III solution contained (in mM): 140.0 NaCl, 5.4 KCl, 1.8 CaCl₂, 1.0 MgCl₂, 5.0 HEPES, and 5.5 glucose, adjusted to pH 7.4 with NaOH. The internal pipette solution contained (in mM): 5.0 NaCl, 140.0 KCl, 10.0 HEPES, 5.0 MgATP, and 10.0 EGTA, adjusted to pH 7.2 with NaOH.

#### Experimental protocols

Freshly dissociated SMCs were allowed to settle on glass coverslips for at least 5 minutes before recording. After achieving whole-cell configuration, SMCs were held at −50 mV and dialyzed with internal solution for at least 2 minutes before starting protocols. Solution changes were achieved using a gravity-fed perfusion system. Patch-clamp experiments were performed at room temperature (22–24°C).

##### Yoda2 protocols

For agonist-only experiments, a 30-second baseline was recorded in 1.8 mM Ca²⁺ Tyrode’s III, followed by 90 seconds of Yoda2 (10µM) in 1.8 mM Ca²⁺ Tyrode’s III and then washout with drug-free 1.8 mM Ca²⁺ Tyrode’s III. For Yoda2 plus paxilline experiments, a 30-second baseline was followed by 90 seconds of Yoda2 (10 µM), 90 seconds of Yoda2 plus paxilline (10 µM), and then washout with 1.8 mM Ca²⁺ Tyrode’s III.

##### Thapsigargin protocol

For thapsigargin (TG) experiments, ISMCs were preincubated for 20 minutes in 1.8 mM Ca²⁺ Tyrode’s III containing TG (1µM) and allowed to adhere to glass coverslips. After establishing whole-cell configuration at -50mV, cells were continuously perfused with 1.8 mM Ca²⁺ Tyrode’s III containing 1 µM TG; following a 30-second baseline, perfusion was switched to the same solution containing Yoda2 (10µM) for 90 seconds and then returned to thapsigargin-only solution for a 30-second washout.

Acute Ca²⁺ removal protocol. To examine acute extracellular Ca^2+^ removal, a 30-second baseline was recorded in 1.8 mM Ca^2+^ Tyrode’s III. The solution was then switched to Ca^2+^-free Tyrode’s III for 30 seconds. Cells were perfused for 90 seconds with 0 mM Ca^2+^ Tyrode’s III containing Yoda2 (10 µM), then for 90 seconds with Yoda2 plus paxilline (10 µM), followed by a Ca^2+^-free washout, and finally re-exposure to 1.8 mM Ca^2+^ Tyrode’s III for 30 seconds.

#### Analysis

Patch clamp data was exported from Clampfit software (Molecular Devices, Sunnyvale, CA) and analyzed in Microsoft Excel. Means to describe various drug conditions were obtained by averaging pA values during the last 10 seconds of the baseline, thapsigargin, and paxilline conditions. For the Yoda2 condition, pA values at the apex of the curve were averaged. Values were normalized to the cell capacitance in order to account for physiological variation across cell sizes

### Immunocytochemistry/Immunofluorescence

#### Immunostaining

SMCs were fixed in 4% paraformaldehyde, quenched with 50 mM glycine, and stained with Alexa Fluor 488-conjugated WGA (5 µg/µl). Cells were blocked in 3% BSA and 0.25% Triton X-100 before overnight incubation with anti-RyR2 (1:50) and anti-RFP (1:100). Secondary antibodies included 568-conjugated goat anti-chicken IgY and Cy5-conjugated goat anti-mouse IgG. Images were captured using a Dragonfly 200 spinning disc confocal (Andor) through a 60x oil immersion objective (NA 1.40). Z-stacks were acquired with 178 nm separation and processed using 3D deconvolution.

#### Imaging Acquisition

Immunofluorescence images were captured using a Dragonfly 200 spinning disc confocal (Andor) through a DMi* Leica microscope (Leica) equipped with a 60x oil immersion objective (NA 1.40) and Andor iXon EMCCD cameras. Laser lines at 488, 561, and 637 nm were utilized to specifically excite WGA-488, RFP/Alexa Fluor-568, and Alexa Fluor-647, respectively. Image acquisition was conducted using Fusion software (Andor) to capture z-stacks with a separation of 178 nm between image slices and apply 3-dimensional deconvolution to image stacks.

#### Immunofluorescence Image Analysis

Image stacks were segmented in Imaris 10.2.0 software (Oxford Instruments). WGA-488 signal was traced manually to delineate the plasma membrane using the Surfaces tool. RyR and Piezo1-associated fluorescence signals were mapped to x/y/z centroid coordinates using the Spots tool, assigning Spots to any fluorescence signals exceeding a fixed threshold with a diameter exceeding 120 nm in all dimensions. ‘Region Growing’ was then applied to assign variable sizes to Spots, determined by the size and brightness of their corresponding fluorescent puncta. Spots were assigned to plasma membrane associated (Membrane) and non-membrane associated (Internal) groups, delimited by a 200 nm distance between the centroid x/y/z position of the Spots and the plasma membrane Surface, and colocalization of Piezo1 with RyR clusters was determined by a 240 nm centroid coordinate proximity cutoff.

### Proximity Ligation Assay

Proximity ligation assays (PLA) were performed using the Duolink PLA system previously described (Martín-Aragón Baudel *et al*. (2022); Sigma, #DUO92008) to detect Piezo1-containing complexes with either ryanodine receptors (RyR) or BKα subunits. Freshly dissociated smooth muscle cells were plated onto glass coverslips and allowed to adhere. Cells were fixed in glyoxal solution (Sigma-Aldrich, #128465) for 20 min, washed in PBS, and quenched with 100 mM glycine (Sigma-Aldrich, #50046) in PBS for 15 min, followed by two additional 5-min washes in PBS. Cells were permeabilized with 0.1% Triton X-100 for 20 min and blocked in 20% Intercept Blocking Buffer (LI-COR Biosciences, #927-60001) for 1 h at 37 °C. Cells were incubated overnight at 4 °C with primary antibodies against Piezo1 (rabbit anti-Piezo1; Alomone APC-087, 1:200) in combination with either mouse anti-RyR2 (Invitrogen C3-33, 1:200) or mouse anti-BKα (NeuroMab clone L6/60, 1:200). After primary antibody incubation, cells were washed three times for 10 min in PBS. PLA detection was performed using Duolink rabbit PLUS and mouse MINUS probes (1:5 dilution in Duolink Antibody Diluent) for 1 h at 37 °C. Cells were washed three times for 5 min in Duolink Buffer A and incubated with ligation solution for 30 min at 37 °C to allow hybridization and circular DNA template formation at sites of dual labeling. Following three additional 3-min washes in Buffer A, cells were incubated with amplification solution for 120 min at 37 °C. Coverslips were then washed twice for 10 min in Duolink Buffer B, followed by a final 1-min wash in 1% Buffer B diluted in dH₂O. Coverslips were air-dried and mounted using Duolink mounting medium (#DUO82040).

PLA signals were imaged using an Olympus Fluoview FV3000 confocal microscope equipped with a 60× oil-immersion objective (NA 1.42) and OBIS 405- and 561-nm lasers, controlled with Olympus 31S-SW software. Confocal z-stacks were acquired at 0.5-µm step size using constant acquisition settings across conditions. Maximum-intensity projections were generated using Fiji (v2.17.0). PLA puncta were quantified in the 561-nm channel using a custom Fiji macro with a fixed threshold applied uniformly across all images.)

### Statistical Analysis

Data are presented as mean ± SD. Normality was assessed using the Shapiro-Wilk test (all p > 0.05). Paired comparisons within the same preparation or cell were analyzed using paired t-tests. Multiple treatment conditions were analyzed using repeated measures one-way ANOVA with Geisser-Greenhouse correction and Bonferroni post-hoc tests. To account for pseudo-replication, technical replicates were averaged within each animal before statistical comparison, ensuring biological variation (N = animals) determined statistical power. Sample sizes are reported as n (cells/segments/preparations) from N (animals). Sex-specific differences were assessed using two-way ANOVA (sex × treatment). No significant sex × treatment interactions were detected (all p > 0.18), justifying pooled analysis. Effect sizes (Cohen’s d) were calculated for all paired comparisons. Statistical significance was set at p < 0.05. All analyses were performed using GraphPad Prism 10 (RRID:SCR_002798). Complete statistical details are provided in **Tables 1 and S1-S5**.

## RESULTS

### Piezo1 activation selectively impairs force generation and contraction kinetics

To investigate how mechanotransduction regulates intestinal contractility, we performed wire myography on adult mouse small intestinal segments using two preparations (**Fig. 1A**): intact full-thickness segments containing all tissue layers (mucosa, submucosa, muscularis externa, and serosa), and isolated muscularis externa preparations where the mucosa and submucosa were carefully removed while maintaining both smooth muscle layers and the myenteric plexus. Both preparations were mounted in organ baths containing oxygenated Krebs solution at 37 °C under physiological preload conditions that support spontaneous phasic contractions. From these contractions, we quantified steady-state control parameters (**Fig. 1B)** across several biomechanical domains. We recorded amplitude (peak force/contractile strength), integral force (cumulative mechanical work), frequency (contractions per minute), and period coefficient of variation (CV = SD/mean), a reflection of the temporal regularity of contraction spacing and duration.

**FIGURE 1.**
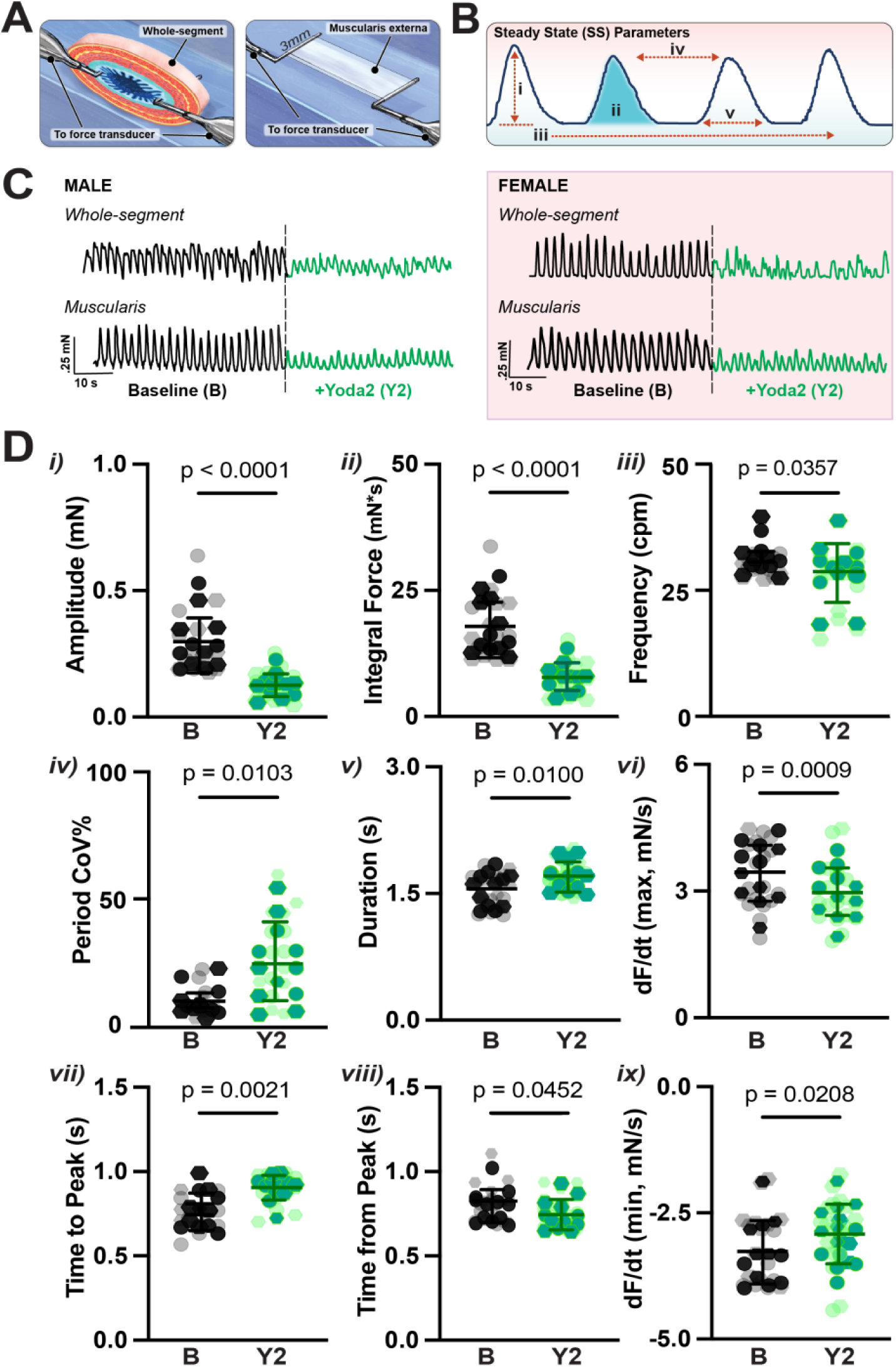
Yoda2 suppresses spontaneous phasic contractions in mouse intestinal smooth muscle. **(A)** Schematic diagrams illustrating wire myography experimental preparations. Left: Whole-segment preparation containing all tissue layers (mucosa, submucosa, muscularis externa, serosa) to preserve physiological tissue architecture. Right: Isolated muscularis externa preparation with mucosa and submucosa removed to eliminate non-muscle cell populations. **(B)** Primary contractile parameters depicted on schematic with the following definitions: (**i**) *amplitude*, peak force generated during each contraction cycle; **(ii)** *integral force*, total area under all contractions representing cumulative mechanical work; **(iii)** *frequency*, number of contractions per minute; **(iv)** *period coefficient of variation (CV = SD/mean)*, temporal regularity of contraction spacing; **(v)** *duration,* time from contraction onset to return to baseline. **(C)** Representative traces showing spontaneous phasic contractions at baseline (black) and after Yoda2 (25 μM, green) in whole-segment (top) and isolated muscularis (bottom) preparations from male (left) and female (right) mice. Scale bars: 0.25 mN, 10 s. **(D)** Contractile parameter analysis comparing baseline (black) vs. Yoda2 (green). Males: circles; females: hexagons. Opaque symbols: per-animal averages; translucent: individual replicates. (**i–iii**) Yoda2 reduced amplitude (57%), integral force (56%), and frequency (9%). (**iv–v**) Temporal regularity decreased (period CoV increased 152%; duration increased 10%). (**vi–ix**) Contraction velocity decreased (dF/dt max reduced 14%; time to peak increased 17%) while relaxation remained proportionally intact (dF/dt min reduced 11% vs. 57% amplitude decline). n = 18 preparations from N = 12 mice (5M/7F). Data: mean ± SD. Statistics: paired t-tests on animal-averaged values. See **Table 1, S1** for complete statistics.

**Table 1.**
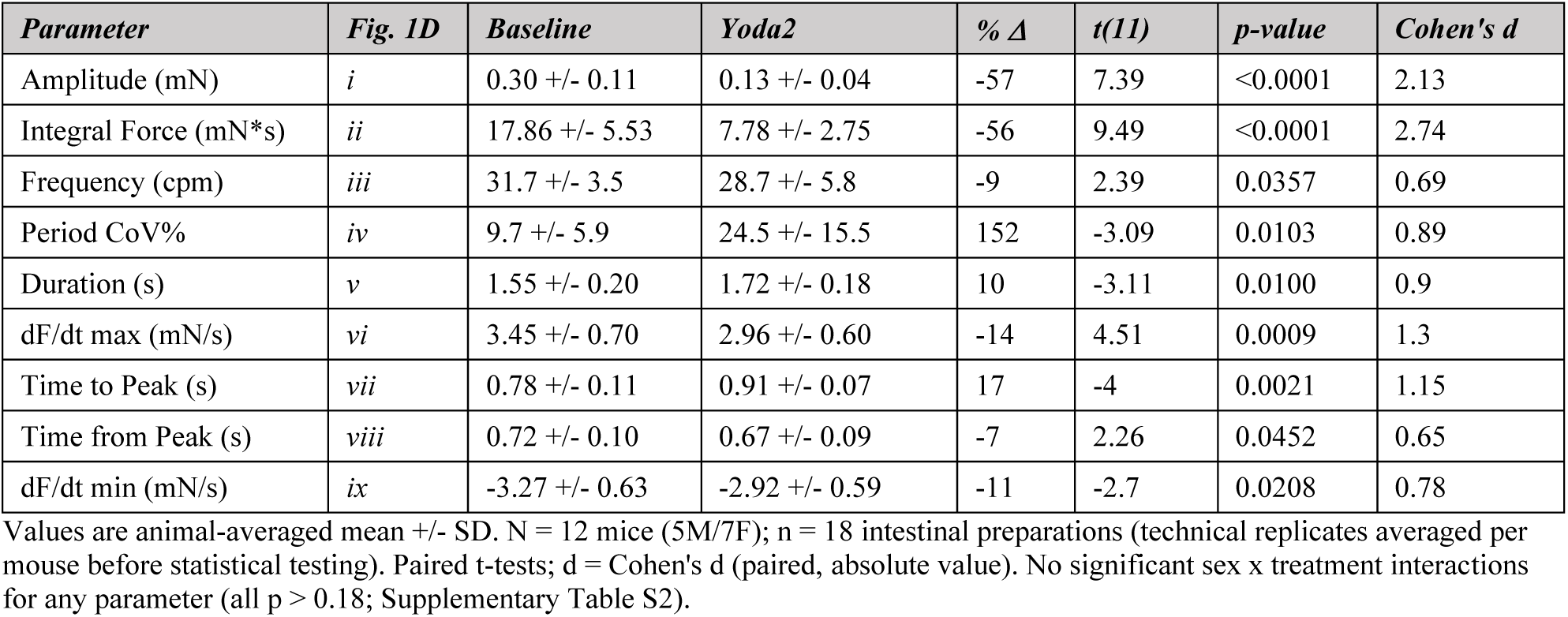
Steady-state contractile parameters before and after Piezo1 activation with Yoda2.

| *Parameter* | *Fig. 1D* | *Baseline* | *Yoda2* | *% Δ* | *t(11)* | *p-value* | *Cohen's d* |
| --- | --- | --- | --- | --- | --- | --- | --- |
| Amplitude (mN) | *i* | 0.30 +/- 0.11 | 0.13 +/- 0.04 | -57 | 7.39 | <0.0001 | 2.13 |
| Integral Force (mN*s) | *ii* | 17.86 +/- 5.53 | 7.78 +/- 2.75 | -56 | 9.49 | <0.0001 | 2.74 |
| Frequency (cpm) | *iii* | 31.7 +/- 3.5 | 28.7 +/- 5.8 | -9 | 2.39 | 0.0357 | 0.69 |
| Period CoV% | *iv* | 9.7 +/- 5.9 | 24.5 +/- 15.5 | 152 | -3.09 | 0.0103 | 0.89 |
| Duration (s) | *v* | 1.55 +/- 0.20 | 1.72 +/- 0.18 | 10 | -3.11 | 0.0100 | 0.9 |
| dF/dt max (mN/s) | *vi* | 3.45 +/- 0.70 | 2.96 +/- 0.60 | -14 | 4.51 | 0.0009 | 1.3 |
| Time to Peak (s) | *vii* | 0.78 +/- 0.11 | 0.91 +/- 0.07 | 17 | -4 | 0.0021 | 1.15 |
| Time from Peak (s) | *viii* | 0.72 +/- 0.10 | 0.67 +/- 0.09 | -7 | 2.26 | 0.0452 | 0.65 |
| dF/dt min (mN/s) | *ix* | -3.27 +/- 0.63 | -2.92 +/- 0.59 | -11 | -2.7 | 0.0208 | 0.78 |

Activation of Piezo1 with the selective agonist Yoda2 (25 μM; Qiao *et al*. (2025)) produced marked contractile dysfunction. Representative traces from male and female mice (**Fig. 1C**) show visibly reduced contraction amplitude following Yoda2 application, with contractions persisting at a similar frequency but diminished force. Responses appeared qualitatively similar between sexes, and exploratory analyses confirmed no significant sex x treatment interactions for any of the nine contractile parameters (all p > 0.18; **Supplementary Table S1**). Data from male and female mice were therefore pooled for all subsequent analyses. Because full-thickness intestinal preparations introduce variability from mucosal and submucosal cell populations that can influence contractile measurements independent of smooth muscle function, we used isolated muscularis externa preparations for all subsequent pharmacological experiments.

Paired t-test analysis of the muscularis externa data (**Fig. 1D**; N = 12 mice, 5M/7F; full statistics in **Table 1**) revealed dysfunction across all three biomechanical domains. Peak contraction amplitude decreased 57% (p < 0.0001; **Fig. 1Di**), and integral force decreased 56% (p < 0.0001; **Fig. 1Dii**), indicating substantially diminished capacity to generate the sustained mechanical energy required for normal intestinal motility (Sanders *et al*., 2012). In contrast to the large reductions in force, contraction frequency decreased by just 9% (p = 0.0357; **Fig. 1Diii**) and contraction duration increased by 10% (p = 0.0100; **Fig. 1Dv**), indicating that pacemaker-driven contraction events persist with only modest changes in their temporal profile. However, the period CoV% increased by 152% (p = 0.0103; **Fig. 1Div**), revealing that while the pacemaker rhythm is maintained, the coupling between pacemaker signals and smooth muscle force output becomes less precise (Sanders, 2019).

The modest increase in contraction duration raises the question of whether Yoda2 slows the contractile cycle uniformly or selectively affects specific phases. Waveform kinetics revealed a striking asymmetry between the contraction and relaxation phases (**Fig. 1Dvi-ix**). Peak contraction velocity (dF/dt max) decreased 14% (p = 0.0009; **Fig. 1Dvi**) and time to peak increased 17% (p = 0.0021; **Fig. 1Dvii**), indicating that the force-generating upstroke is substantially slowed. In contrast, the relaxation phase was preserved or relatively enhanced: time from peak decreased 7% (p = 0.0452; **Fig. 1Dviii**), and although the absolute peak relaxation velocity (dF/dt min) decreased 11% in magnitude (p = 0.0208; **Fig. 1Dix**), this small change accompanies a 57% reduction in amplitude, indicating that the tissue relaxes proportionally faster relative to the force it generates.

Together, these data demonstrate that Piezo1 activation profoundly inhibits force generation and selectively slows the contraction phase relative to relaxation, raising the possibility that Yoda2 acts through intracellular Ca^2+^ handling pathways. To test this, we next examined whether depleting SR Ca^2+^ stores or disrupting SR Ca^2+^ release reduces the contractile effects of Yoda2.

### Piezo1-mediated contractile inhibition depends on SR Ca^2+^

We used two complementary approaches to disrupt SR Ca^2+^ signaling prior to Yoda2 application (**Fig. 2**). Thapsigargin irreversibly inhibits SERCA, passively depleting SR Ca^2+^ stores (Lytton *et al*., 1991), while ryanodine locks RyRs in a sub-conductance state, preventing physiological Ca^2+^ release (Rousseau *et al*., 1987). If Yoda2 acts through either of these SR Ca^2+^ pathways, pre-treatment with either drug should reduce its contractile effects.

**FIGURE 2.**
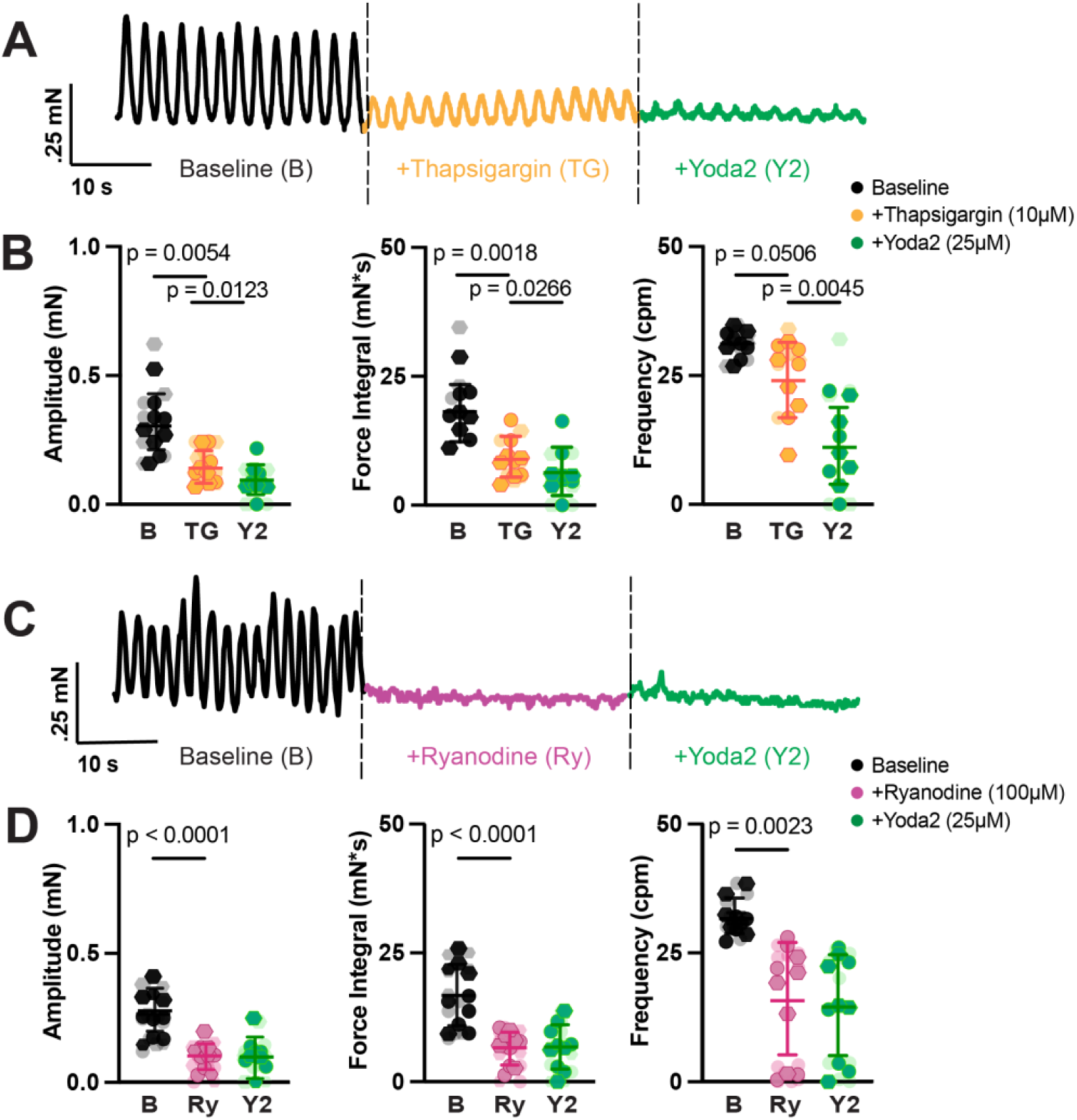
SR Ca^2+^ store disruption attenuates Piezo1-mediated contractile inhibition. (A,B) Thapsigargin. **(A)** Representative traces showing baseline (black), thapsigargin (10 μM), and Yoda2 (25 μM, green) effects. Scale bars: 0.25 mN, 60 s. **(B)** Thapsigargin reduced amplitude, integral force, and frequency; subsequent Yoda2 produced further modest reductions. n = 11 segments from N = 9 mice (4M/5F). **(C,D) Ryanodine. (C)** Representative traces showing baseline (black), ryanodine (100 μM, pink), and Yoda2 (25 μM, green). Scale bars: 0.25 mN, 60 s. **(D)** Ryanodine profoundly reduced all parameters; Yoda2 produced no additional effect. n = 14 segments from N = 10 mice (5M/5F). Translucent symbols: individual preparations; opaque: animal averages. Data: mean ± SD. Statistics: repeated measures one-way ANOVA with Geisser-Greenhouse correction, Bonferroni post-hoc. P-values > 0.05 are considered significant. See **Table S3** for complete statistics. B, baseline; TG, thapsigargin; Ry, ryanodine; Y2, Yoda2.

Thapsigargin (10μM), which depletes SR Ca^2+^ stores by irreversibly inhibiting SERCA, reduced contraction amplitude by 54% (p = 0.0054) and integral force by 51% (p = 0.0018; N = 9 mice, 4M/5F). Frequency was not significantly reduced (23% decrease; p = 0.0506; **Fig. 2A-B**; **Supp. Table 1**), indicating that pacemaker-driven contraction initiation is less dependent on SR Ca^2+^ store content than the contractile machinery itself. Subsequent Yoda2 application to thapsigargin-treated tissue produced additional reductions in amplitude (p = 0.0123), integral force (p = 0.0266), and frequency (p = 0.0045), demonstrating partial but incomplete occlusion of the Yoda2 response or contributions from non-SR sources such as plasma membrane Ca^2+^ entry.

Ryanodine at higher doses (100 μM), prevents physiological RyR-mediated Ca^2+^ release, and produced a more uniform disruption, reducing amplitude by 61% (p < 0.0001), integral force by 61% (p < 0.0001), and frequency by 50% (p = 0.0023; N = 10 mice, 5M/5F; **Fig. 2C-D**). The frequency reduction following ryanodine, in contrast to the preserved frequency after thapsigargin, is consistent with RyR-mediated Ca2+ release contributing to the translation of pacemaker signals into coordinated smooth muscle contractions. Following ryanodine pre-treatment, subsequent Yoda2 application produced no further change in any metric (all p > 0.88; **Supp. Table 1).** However, because ryanodine reduced baseline contractility more severely than thapsigargin, a floor effect in which insufficient contractile reserve remained to detect additional inhibition cannot be fully excluded.

Overall, this data suggests that yoda2 and ryanodine may act through overlapping RyR-dependent pathways, suggesting a framework in which Piezo1 triggers SR Ca²⁺ release via ryanodine receptors, thereby “amplifying” the [Ca^2+^]_i_ to collectively activate BK_Ca_ (Santana *et al*., 1996; Dopico *et al*., 2018). However, the variable effects of ryanodine probably stem from its complex, concentration-dependent actions, and other explanations, such as ryanodine interfering with pacemaker mechanisms independently of Piezo1, are also possible (Collier *et al*., 2000).

Regardless, while this data provides compelling evidence that Yoda2-mediated contractile inhibition depends on functional SR Ca^2+^ stores and RyR-mediated release. This would support the hypothesis that Piezo1 mobilizes SR Ca²⁺ to activate hyperpolarizing conductances that inhibit contractility. However, wire myography cannot resolve the underlying cellular mechanism, whether Piezo1 directly triggers SR Ca^2+^ release, activates downstream Ca^2+^ -sensitive conductances, or both. To address this, we performed whole-cell patch-clamp recordings on isolated smooth muscle cells to determine whether Piezo1 activation produces identifiable ionic currents.

### Piezo1 activation produces BK channel-mediated outward currents in isolated smooth muscle cells

To investigate the cellular mechanisms underlying Piezo1-mediated contractile inhibition, we performed whole-cell patch-clamp recordings on freshly isolated smooth muscle cells to determine whether Piezo1 activation produces outward (hyperpolarizing) currents that would lead to relaxation in GI smooth muscle.

At -50 mV, plasma membrane Piezo1 activation would be expected to produce inward (negative) current from Ca²⁺ and Na⁺ influx (Coste *et al*., 2010; Gnanasambandam *et al*., 2015), which would depolarize the membrane and potentially trigger voltage-gated Ca²⁺ channels to enhance contraction. Instead, application of the Piezo1 agonist Yoda2 (10 µM) produced marked outward (positive) currents that developed over 30-60 seconds and returned to baseline upon washout (**Fig. 3A**). Addition of Yoda2 significantly increased current density from -0.30 ± 0.36 pA/pF at baseline to 2.49 ± 1.81 pA/pF during Yoda2 application (p = 0.0015; n = 18 cells from N = 7 mice; **Fig. 3B**), The outward direction indicates K⁺ efflux, which would hyperpolarize the membrane and reduce voltage-gated Ca²⁺ channel activity, consistent with the contractile inhibition observed in tissue recordings.

**FIGURE 3.**
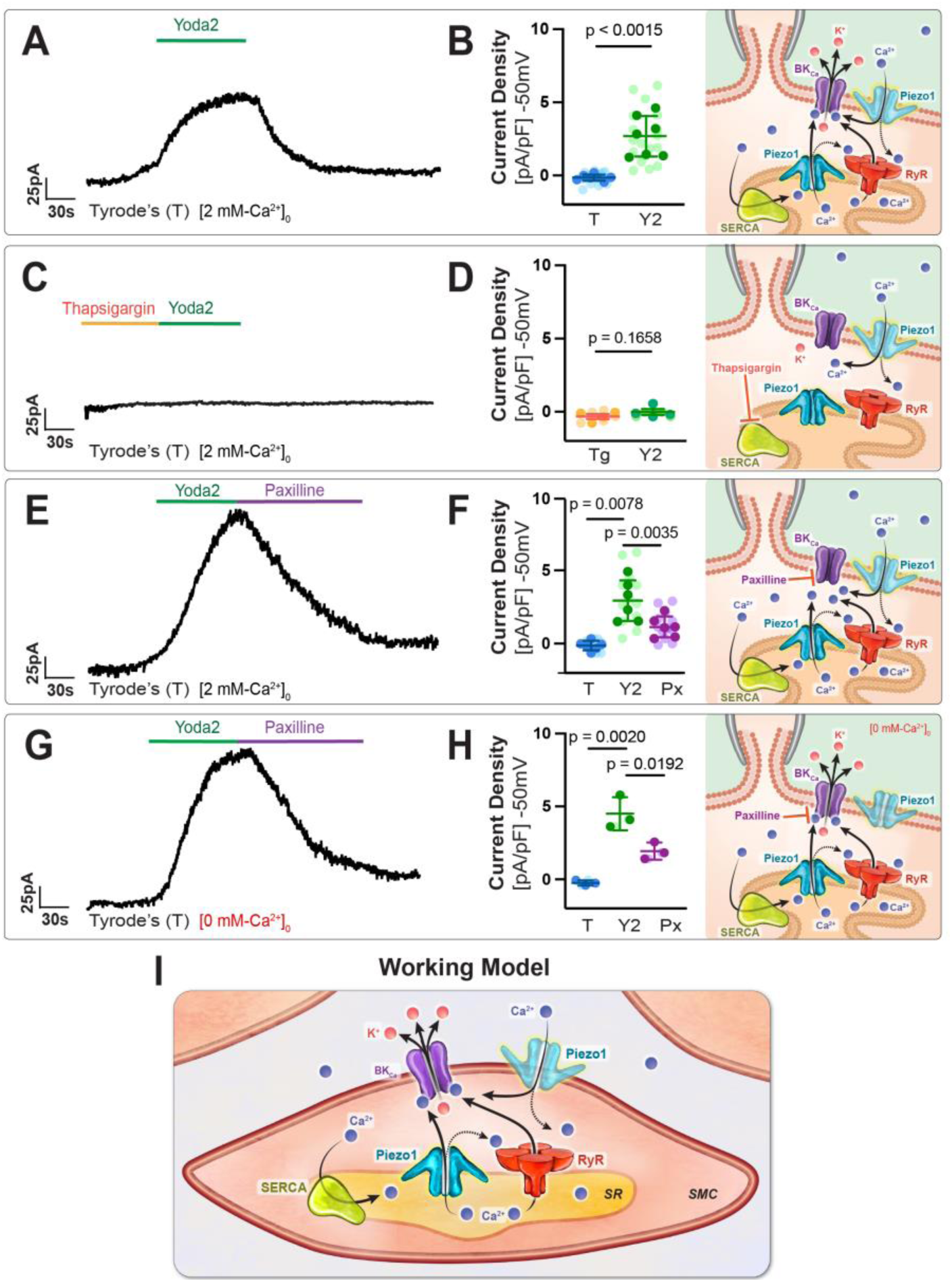
Piezo1 activation produces BK channel-mediated outward currents that require SR Ca²⁺ and persist in 0 mM extracellular Ca²⁺. A-B *Yoda2 evokes outward currents in intestinal smooth muscle cells (iSMCs).* (A) Representative whole-cell patch-clamp current recording from freshly dissociated iSMC voltage-clamped at -50 mV in Tyrode’s III solution at baseline, in response to yoda2 (10 μM, green bar), and after washout. **(B)** Current density (pA/pF) at baseline (T) and peak yoda2-evoked currents (Y2). Paired t-test; N=7 mice, n = 18 cells. The schematic depicts the proposed signaling pathway: Piezo1-mediated Ca^2+^ entry and/or intracellular store release activate ryanodine receptors (RyR) and/or directly activate BK_Ca_ channels. C-D *SR Ca^2+^ store depletion attenuates the Yoda2 response*. **(C)** Representative recording from a cell pre-incubated with thapsigargin (TG) and voltage-clamped at -50 mV at baseline, in response to Yoda2 and washout. **(D)** Current density at baseline with TG during Yoda2. Paired t-test; N = 5 mice, n = 7 cells. Schematic illustrates that SERCA inhibition with TG results in depletion of intracellular Ca^2+^ necessary for Yoda-evoked Piezo1 activation of outward current. E-F *BK channel blockade reduces Yoda2-evoked currents in 2 mM Ca^2^*^+.^ (E) Representative currents recorded at baseline, with Yoda2 (10 μM, green bar) and subsequent addition of paxilline (10 μM, purple bar). **(F)** Current density at baseline (T), with Yoda2 (Y2) and with Paxilline (Px). RM one-way ANOVA; N = 6 mice, n = 10 cells. Schematic illustrating that paxilline inhibits downstream BKCa channels while the Piezo1-RyR Ca^2+^ release pathway remains intact, reducing outward current density. G-H *Yoda2-evoked, paxilline-sensitive currents persist without extracellular Ca^2+^* (G). Representative recording at baseline, after acute removal of extracellular Ca^2+^, during Yoda2 (green bar), after paxilline (purple bar), and washout. **(H)** Current density at baseline in 0 mM Ca2+ (T), peak Yoda2 (Y2), and after paxilline (Px). N = 3 mice. Schematic illustrates that [Ca^2+^]_i_ store released through RyR sustains BKCa activation in the absence of extracellular Ca^2+^ entry, and paxilline reduces the outward current. Individual cells are shown as translucent dots with opaque dots for per-animal averages; data shown as mean ± SD. P-values as indicated above with p<0.05 marked as significant. **(I)** Working model illustrating Piezo1-RyR-BK_Ca_ axis. Activation of Piezo1 in the plasma membrane and SR increase cytosolic Ca^2+^ triggering additional Ca^2+^ release from intracellular stores via RyR. The combine Ca^2+^ signals locally activate nearby BK_Ca_ channels resulting in K+ efflux and membrane hyperpolarization while SERCA maintains SR Ca^2+^ content to sustain the signaling loop.

To test whether the Yoda2-evoked outward current requires SR Ca²⁺ stores, cells were pre-incubated with thapsigargin (1 µM, 20 min) to impair SR Ca²⁺ uptake and deplete releasable SR Ca²⁺ before recording. Following thapsigargin pretreatment, cells were perfused continuously with thapsigargin-containing solution to maintain store depletion throughout the recording (**Fig. 3C**). Under these conditions, Yoda2 application did not significantly alter current density (p = 0.1656; n=7 cells from 5 mice; **Fig. 3D**), demonstrating that functional SR Ca²⁺ stores are necessary for Piezo1-mediated outward currents.

The outward current kinetics and Ca²⁺ dependence are consistent with BK channel activation, which is driven by local Ca²⁺ release events (Ca²⁺ sparks) in smooth muscle (Nelson et al., 1995; Jaggar et al., 1998). We therefore tested this by applying the selective BK channel blocker paxilline (10 µM) during the Yoda2 response, which significantly reduced the Yoda2-evoked current from 2.83 ± 0.57 to 1.04 ± 0.29 pA/pF (p = 0.0035, N=6 mice, n=10).

To determine whether the Yoda2-induced current requires Ca²⁺ influx from the extracellular space, we acutely removed extracellular Ca²⁺ sources by switching to Ca²⁺-free Tyrode’s II after establishing an initial 30s baseline. In 0 [Ca²⁺]o external solution, Yoda2 still produced robust outward currents (4.82 ± 0.683 pA/pF vs. -0.11 ± 0.109 pA/pF at baseline; p = 0.0020; N = 3 mice; Fig. 3H). Subsequent application of paxilline (10 µM) reduced current density to 2.15 ± 0.3531 pA/pF (p = 0.0192), representing 54% inhibition of the Yoda2-induced increase. The persistence of Yoda2-evoked, paxilline-sensitive outward currents under Ca^2+-^free conditions suggests that extracellular Ca^2+^ entry is not the main trigger. Instead, it points to intracellular Ca^2+^ release as the source for activating BK channels. However, the possibility that residual extracellular Ca^2+^ or Ca^2+^ that entered prior to the solution change cannot be entirely ruled out.

Together, these electrophysiological findings reveal that Piezo1 activation in small intestinal smooth muscle cells produces outward K⁺ currents that would be expected to cause membrane hyperpolarization and contractile inhibition. The BK channel-mediated outward currents suggest involvement of SR Ca²⁺ signaling, as BK channel activation in smooth muscle typically occurs through Ca²⁺ spark-mediated local Ca²⁺ elevations. To determine whether Piezo1, RyR, and BK channels are positioned close enough to support this local signaling mechanism, we next examined their spatial relationships using immunofluorescence and proximity ligation assays.

### Piezo1 predominantly localizes to intracellular membranes in intestinal smooth muscle cells

To determine whether the functional coupling between Piezo1, SR Ca²⁺ release, and BK channel activation observed (**Fig.1-3**) reflects specific subcellular organization, we examined the spatial distribution of Piezo1, RyR, and BKCa in freshly isolated intestinal smooth muscle cells from Piezo1-tdTomato reporter mice. Piezo1 localization was detected using anti-RFP antibodies against the TdTomato reporter, which provided more reliable targeting than direct Piezo1 antibodies, given challenges with Piezo1 immunolocalization (Alper, 2017).

Immunocytochemical labeling (**Fig. 4A**) revealed that Piezo1 clusters (anti-RFP, cyan) were predominantly distributed within intracellular compartments rather than at the plasma membrane, following a pattern similar to that of RyR (magenta; **Fig. 4A**). Quantitative analysis performed using surface-based classification algorithms (Imaris, ±2 pixel threshold relative to membrane surface) on N=10 mice, 5M/5F demonstrated that 86.15 ± 7.28% of Piezo1 clusters were intracellular (cyan) compared to 13.85 ± 7.28% at the plasma membrane (blue; p<0.0001; **Fig. 4B, E**). Similarly, RyR immunoreactivity was detected predominantly in intracellular compartments (magenta) rather than near the plasma membrane (red; p<0.0001; **Fig. 4C,F**). This predominant intracellular distribution of both Piezo1 and RyR is consistent with SR localization and with the functional requirement for SR Ca2+ stores, as demonstrated by thapsigargin sensitivity (Fig. 2). No significant sex differences were detected for any parameter (all interaction p > 0.35; **supp Table 1**).

**FIGURE 4:**
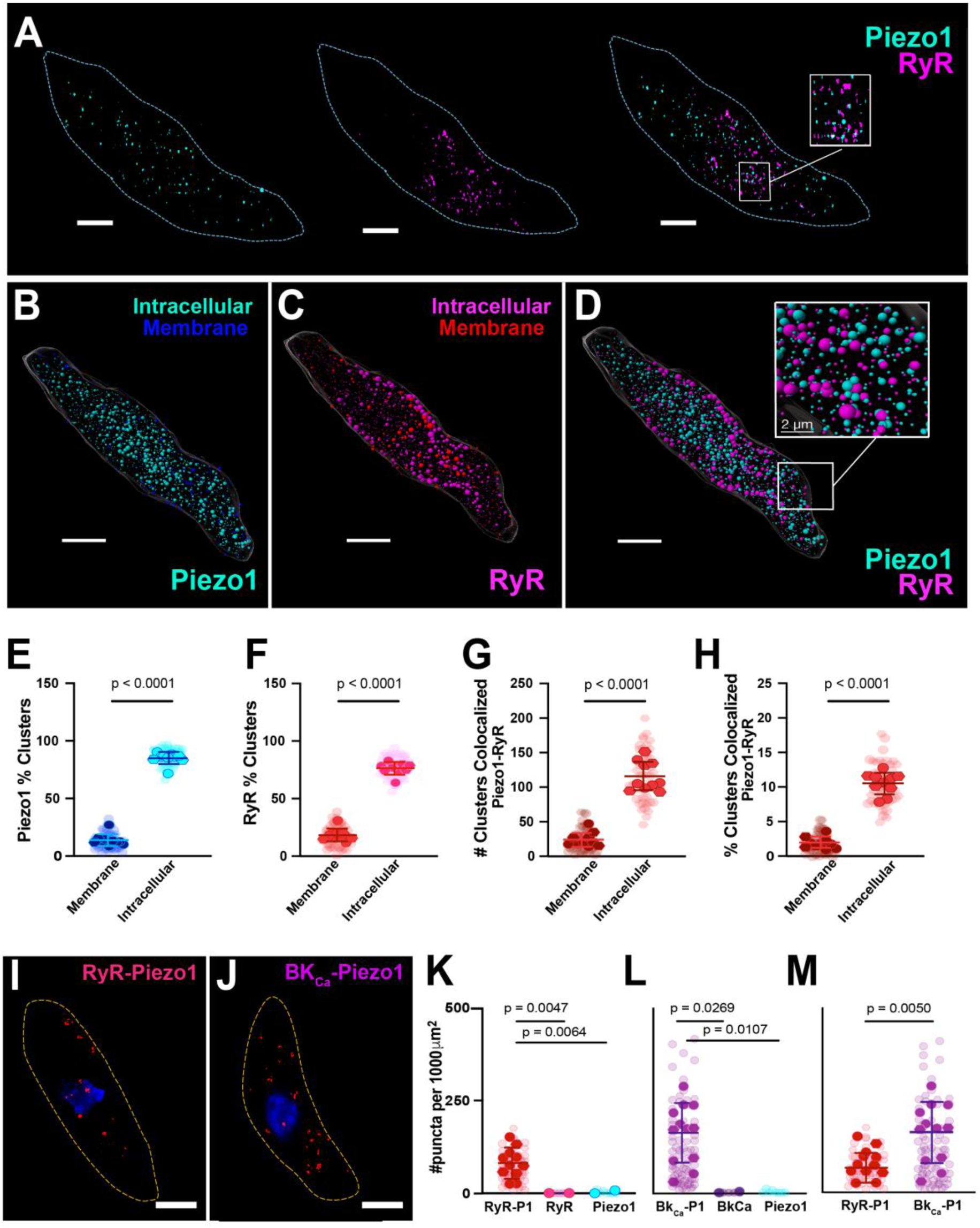
Piezo1 localizes predominantly intracellularly with nanoscale proximity to RyR and BK channels. (A–D) Confocal microscopy of jejunal circular smooth muscle from Piezo1-tdTomato mice. (A) Left to right: Piezo1-tdTomato (cyan), RyR (magenta), merge with insets showing colocalization. Scale bars: 10 μm; inset 2 μm. (B,C) Imaris 3D surface classification separating membrane-localized (B: Piezo1 blue, RyR red) from intracellular (C: Piezo1 cyan, RyR magenta) signal **(B,C)** Imaris 3D surface classification separating membrane-localized (B: Piezo1 blue, RyR red) from intracellular (C: Piezo1 cyan, RyR2 magenta) signal. **(D)** Merged view showing extensive intracellular colocalization between Piezo1 (cyan) and RyR (magenta). **(E–H)** Quantification of Piezo1/RyR cluster distribution and association for n=50 cells from N=10 animals: Percentage of Piezo1 **(E)** and RyR **(F)** clusters localized at the plasma membrane (± 2 voxels) or intracellular compartments; the number **(G)** and percentage **(H)** of all Piezo1 clusters colocalized with RyR, either at the plasma membrane or intracellularly. Statistics: P-values as indicated above with p<0.05 marked as significant. data analyzed from n = 48 cells from N = 10 mice (5M/5F). No sex differences detected (all P > 0.6). Nested t-tests/nested two-way ANOVA (**Full statistics in Table S4**). Translucent symbols: individual cells; opaque: animal averages. Data: mean ± SD. Statistics: nested one-way ANOVA (K,L) or nested t-test, n = 127 cells from N = 16 mice. **(I,J)** Proximity ligation assay (PLA) detecting nanoscale interactions (<40 nm proximity). Max-intensity projections show PLA puncta (red) with Dapi (blue) **(I)** RyR-Piezo1 and **(J)** BK_Ca_-Piezo1. **(K-M)** Quantification (puncta per mm2) of (K) RyR-Piezo1 with 120.5 +/- 105.30 (L) BK_Ca_-Piezo1 67.57+/-39.85 and (M) a direct comparison between the two. n = 127 cells from N = 16 mice. Translucent symbols: individual cells; opaque: animal averages. Data: mean ± SD. Statistics: nested one-way ANOVA (K,L) or nested t-test (M) accounting for within-animal clustering. Single antibody controls verify PLA specificity.

To determine whether Piezo1 and RyR occupy similar subcellular domains, we quantified clusters exhibiting spatial overlap (center-to-center distance < 240 nm) separately for membrane and intracellular compartments. Colocalized Piezo1-RyR clusters were approximately five-fold more abundant in the intracellular compartment compared to the plasma membrane (nested analysis, p < 0.0001; **Fig. 4G**). This intracellular enrichment was also reflected in cluster volume: 89.22 ± 8.08% of total Piezo1 signal volume was intracellular versus 10.80 ± 8.07% at the membrane (nested analysis, p < 0.0001; **Fig. 4H**). These data indicate that Piezo1 and RyR are not merely co-distributed within the same cellular compartment but preferentially co-assemble within intracellular domains.

While confocal microscopy (resolution limit ∼200 nm) can resolve colocalization at the subcellular level, it cannot determine whether proteins are positioned within the <100 nm range typical of functional Ca²⁺ coupling domains. To address this limitation, we employed proximity ligation assay (PLA), which generates fluorescent puncta only when two target proteins are within <40 nm of each other (Fig. 4I–J). RyR-Piezo1 PLA signal was significantly elevated compared to single-antibody controls (nested one-way ANOVA; p = 0.0064 vs. RyR-only, p = 0.0047 vs. Piezo1-only; Fig. 4K), confirming nanoscale proximity between these two channels. BKCa-Piezo1 PLA signal was similarly elevated compared to controls (p = 0.0269 vs. BKCa-only, p = 0.0107 vs. Piezo1-only; Fig. 4L). Notably, BKCa-Piezo1 proximity was approximately two-fold more abundant than RyR-Piezo1 proximity (p = 0.0050; Fig. 4M), suggesting a stoichiometry in which multiple BK channels surround each Piezo1-RyR release site. This arrangement is consistent with established Ca²⁺ spark-STOC coupling, where multiple BK channels must be activated by a single Ca²⁺ release event to produce a physiologically meaningful hyperpolarization (Brenner *et al*., 2000).

Collectively, these data demonstrate that Piezo1, RyR, and BK channels reside within <40 nm of each other at intracellular sites, providing the nanoscale spatial framework necessary for the SR Ca²⁺ store-dependent, BK-mediated outward currents identified in Figs. 2–3.

## DISCUSSION

This study identifies a functional intracellular mechanotransduction axis in small intestinal smooth muscle, defined by a specialized signaling complex on the SR of the small intestine, comprising intracellular Piezo1 (intra-Piezo1) and RyR, in complex with PM BK_Ca_ channels. Three convergent lines of evidence support this conclusion. First, Piezo1 activation with Yoda2 reduced contractile force and slowed contraction kinetics at the tissue level, consistent with the engagement of a relaxation pathway rather than excitatory surface-channel activity. Second, Piezo1 activation evoked outward currents required intact SR Ca²⁺ stores but not extracellular Ca²⁺ and were abolished by the BK_Ca_ channel inhibitor paxilline, placing the signal origin at the sarcoplasmic reticulum. Third, high-resolution imaging and proximity ligation assays revealed that ∼86% of Piezo1 clusters reside intracellularly, colocalize with RyR, and are positioned within <40 nm of both RyR and BK_Ca_ channels.

The present findings define a novel physiological axis in smooth muscle, providing new functional evidence on how organelle mechanosensation occurs via intracellular Piezo1 (intra-Piezo1). The canonical view posits that Piezo1-mediated mechanotransduction is initiated exclusively at the PM via cation influx (Coste *et al*., 2010; Gudipaty *et al*., 2017). Our work identifies an alternative intracellular pathway for smooth muscle cell mechanosensitivity, mediated by intra-Piezo1. We demonstrate that, in small intestinal smooth muscle, this process is fundamentally important and identify a specialized mechanotransduction intracellular signaling pathway: a “Sensor-Amplifier-Effector” complex located on the SR composed of 3 components: Intra--Piezo1 (Sensor), Ryanodine Receptors (Amplifier), and BK_Ca_ channels (Effector). Activation of intracellular Piezo1 would result in Ca^2+^ release from the SR into the cytosolic surface (Liao *et al*., 2021). The localized Ca²⁺ release would then trigger RYR to activate Ca²⁺-induced Ca²⁺ release (CICR) at other nearby RyRs. The resulting amplified Ca^2+^ pool would then activate BK_Ca_ channels, leading to membrane hyperpolarization and muscle relaxation (Nelson *et al*., 1995; Ríos, 2018).

The proposed intracellular axis functions as a molecular brake, coupling organellar mechanosensing via intra-Piezo1 to BK_Ca_-mediated hyperpolarization to dampen excitability and promote relaxation. This resolves the paradox of why knocking out Piezo1 channels in intestinal smooth muscle cells leads not only to contractile deficits but also to complex ion-channel remodeling that creates a hyperexcitable environment (Bautista *et al*., 2025a). Such profound, indirect consequences establish that organelles are active participants in detecting and regulating stress from the inside out.

### Intracellular Piezo1: Evidence for a Subcellular Signaling Hub

The predominantly intracellular localization of Piezo1 in intestinal smooth muscle is not an isolated finding. Growing evidence demonstrates that Piezo1 functions within intracellular compartments across diverse cell types, suggesting this dual localization reflects a conserved property of the channel (Bautista *et al*., 2025b). Rather than simply a surface channel that is incidentally present at intracellular membranes, the emerging picture is of a subcellular signaling hub whose output is determined by its compartmental address and local nanodomain partners.

The most direct smooth muscle precedent comes from pulmonary arterial smooth muscle cells (PASMCs), where Liao *et al*. (2021) Piezo1 colocalization with ER/SR, mitochondrial, and nuclear envelope markers was demonstrated, with Yoda1-evoked Ca²⁺ transients persisting in Ca²⁺-free medium and abolished by Piezo1 siRNA knockdown. In epithelial cells, McHugh *et al*. (2010) used subcellular fractionation and ER marker co-enrichment to localize Fam38A/Piezo1 to the endoplasmic reticulum in HeLa and CHO cells, where it recruits R-Ras, increases Ca²⁺ release from cytoplasmic stores to activate calpain, and triggers β1 integrin activation through talin cleavage, an inside-out signaling pathway with consequences for cell adhesion and migration (McHugh *et al*., 2010; McHugh *et al*., 2012). This diversity of locations and functional outputs, ranging from definitive organellar residence to functional coupling, argues against a trafficking artifact, as biosynthetic intermediates would not produce such organelle-appropriate signaling across cell types.

Piezo1’s biophysical properties rationalize how the channel operates at intracellular membranes. Its large structural footprint and curvature sensitivity favor low-curvature membranes such as the ER/SR, and its capacity for conformational signaling, mechanical rearrangement of propeller domains independent of ion flux, enables mechanotransduction at organellar membranes with ionic compositions distinct from the extracellular space (Guo & MacKinnon, 2017; Lewis *et al*., 2024). Critically, the ER/SR membrane is not mechanically isolated: it experiences direct tension as part of a coupled membrane continuum, with PM curvature regulating ER contact formation (Yang *et al*., 2024; Townson & Progida, 2025). In endothelial cells, Piezo1 promotes ER Ca²⁺ transport in response to shear stress through an IP_3_R2-dependent circuit (Santana Nunez *et al*., 2023), and in the Drosophila heart, Piezo buffers mechanical stress through intracellular Ca²⁺ modulation rather than surface currents (Zechini *et al*., 2022). Together, these findings establish that mechanical force can reach and gate intracellular Piezo1 across organ systems and species (Bautista *et al*., 2025b).

A unifying principle emerges: Piezo1 operates as a mechanosensory hub, with its subcellular address and nanodomain partners dictating the direction and magnitude of its output. This model, in which the same channel produces hyperpolarization from the SR and depolarization from the plasma membrane, resolves the longstanding paradox of how a single ion channel can promote both proliferation and apoptosis, contraction and relaxation (Gudipaty *et al*., 2017).

### The Nanoscale Architecture of the Organellar Mechanosensory Niche

The Sensor-Amplifier-Effector complex we describe is built upon a well-characterized foundation of nanoscale Ca²⁺ signaling at PM-SR junctions, a fundamental organizing principle of smooth muscle physiology. Within the ∼20 nm gap of peripheral junctions, localized Ca²⁺ sparks from RyR in the SR activate clustered BK_Ca_ channels on the overlying PM, producing spontaneous transient outward currents (STOCs) that oppose voltage-gated Ca²⁺ entry and constrain myogenic tone, establishing the paradigm of spatially restricted Ca²⁺ coupling in vascular smooth muscle (Nelson *et al*., 1995; Santana *et al*., 1996; Jaggar *et al*., 2000). This system requires nanometer-scale co-assembly of RyR and BK_Ca_ channels at junctional sites organized by scaffolding proteins such as junctophilin-2 (Santana *et al*., 1996; Kotlikoff, 2003; Pritchard *et al*., 2019).

In the GI tract, BK_Ca_ channels function as essential brakes on excitability (Meredith *et al*., 2006; Wang *et al*., 2010; Ren *et al*., 2016), yet their precise contributions and activation mechanisms vary across intestinal segments. BK_Ca_ channel function is best characterized in the colon, where the inhibitory architecture differs fundamentally from the classical RyR-BK_Ca_ spark-STOC paradigm. In colonic smooth muscle, spontaneous Ca²⁺ transients are IP₃R-mediated Ca²⁺ puffs that couple to spontaneous transient outward currents through both BK_Ca_ and small-conductance SK channels (Koh *et al*., 1998; Bayguinov *et al*., 2000; Bayguinov *et al*., 2001). However, in the small intestine, STOCs are driven instead by RyR-dependent Ca²⁺ sparks that couple directly to BK_Ca_ channels (Benham *et al*., 1986; Bolton & Lim, 1989; Nelson *et al*., 1995), with localized Ca²⁺ release events later visualized directly in guinea-pig ileal myocytes by confocal microscopy (Gordienko *et al*., 1998). These foundational studies established that small intestinal STOCs couple RyR-mediated Ca²⁺ release to BK channel activation (Bolton & Imaizumi, 1996), but left two critical questions unanswered: what triggers spontaneous RyR opening in these cells, and how does the spark-STOC pathway connect to functional motility outcomes?

In vascular smooth muscle, both answers are clear: Ca²⁺ entry through L-type voltage-gated channels refills SR stores and provides the CICR trigger for RyR-mediated sparks, and BK β1-subunits confer sufficient Ca²⁺ sensitivity for spark-STOC coupling to regulate myogenic tone (Jaggar *et al*., 2000; Takeda *et al*., 2011; Dopico *et al*., 2018). However, while the small intestinal RyR-BK axis was established pharmacologically decades ago, the upstream trigger that initiates RyR-dependent Ca²⁺ release remained undefined, suggesting an alternative, potentially mechanosensitive mechanism.

Our study provides the structural basis for the distribution of Piezo1 in small intestinal SMC and its organization with RYRs and BK channels. We defined in this study that intra-Piezo1 and RyR clusters reside intracellularly, with enrichment of intra-Piezo1-RyR colocalization in intracellular domains. Intra-Piezo1 is in proximity to both RyR and BK_Ca_ channels within <40 nm, with a greater abundance of intra-Piezo1-BK_Ca_ puncta, suggesting that multiple BK_Ca_ channels likely surround intra-Piezo1-RyR release sites, recapitulating the amplification logic of vascular spark-STOC coupling while adding a mechanosensitive trigger to the triad (Brenner *et al*., 2000; Plüger *et al*., 2000).

### Physiological Consequences: The Molecular Brake on GI Motility

Piezo1 activation with Yoda2, slowed force muscle development while the relaxation phase was relatively preserved despite an overall 57% reduction in amplitude (**Fig. 1D**). This asymmetry is consistent with BK_Ca_-mediated hyperpolarization reducing voltage-gated Ca²⁺ channel activity (Nelson *et al*., 1995; Hill-Eubanks *et al*., 2011), which would blunt Ca²⁺-dependent myosin light chain phosphorylation and slow force generation, while leaving SERCA-mediated Ca²⁺ reuptake and myosin phosphatase activity, the principal drivers of relaxation, unaffected (Hill-Eubanks *et al*., 2011). The brake induced by Yoda2-mediated Piezo1 activation thus appears to act selectively on excitation-contraction coupling rather than globally suppressing the contractile cycle. The 152% increase in period variability, accompanied by only a small reduction in frequency, may reflect a direct effect on ICC pacemaker activity or an indirect effect mediated by other cell types (Won *et al*., 2005; Sanders *et al*., 2023).

The mechanotransduction brake in smooth muscle cells demonstrated in this study may contribute to intestinal peristaltic coordination. Luminal distension stretches the circular smooth muscle layer, activating SR-resident Piezo1 and triggering Ca²⁺ sparks, BK_Ca_-mediated STOCs, and membrane hyperpolarization, ensuring the muscle segment remains relaxed and preventing excessive wall tension that could obstruct flow. As the bolus advances, decreasing wall tension would reduce Piezo1 activation and release the brake, allowing excitatory inputs from interstitial cells of Cajal and enteric neurons to drive contraction (Sanders *et al*., 2023). If correct, the spatial gradient of mechanical stimulation would create a corresponding gradient of brake engagement: strongly engaged in the distended receiving region, then progressively less engaged upstream, producing the coordinated relaxation-contraction wave characteristic of peristalsis.

This model extends the “mechanical brakes” concept of Joshi et al. Joshi *et al*. (2021), with our data providing a candidate molecular architecture: a Piezo1-RyR-BK_Ca_ complex whose brake strength could scale with junction density and mechanical strain. The prediction is testable: loss of the brake should disrupt coordinated mechano-relaxation, and our previous Piezo1-deficient mice show impaired transit and altered fecal pellet characteristics (Bautista *et al*., 2025a), consistent with, though not definitive proof of, this mechanism. Direct demonstration would require measuring Piezo1-dependent BK_Ca_ currents in response to physiological stretch rather than pharmacological activation. Furthermore, our PLA quantification revealed that the density of the Piezo1-RyR-BK_Ca_ complex varies by more than an order of magnitude across individual cells. This variability suggests these junctions are highly dynamic, actively trafficked by the cytoskeleton in response to mechanical demand, much like the junctional SR motility observed in cardiac myocytes (Drum *et al*., 2020).

### Conclusion

This study fundamentally revises our understanding of mechanotransduction in the GI tract. We demonstrate that Piezo1 is not merely a surface portal for excitation but a dual-function sensor that, when anchored to the sarcoplasmic reticulum, confers mechanosensitivity to intracellular organelles. By recruiting RyR and BK_Ca_ channels into a nanoscale signaling hub, intra-Piezo1 creates a feedback loop that fine-tunes excitability from the inside out, ensuring the robust yet regulated motility essential for life.

## DATA AVAILABILITY

All data supporting the findings of this study are available within the manuscript and its supplementary materials. For experiments with sample sizes n≤30, individual data points are plotted in the figures. Source data for all figures are provided as Supplementary Data files. Raw electrophysiology recordings, confocal imaging datasets, and wire myography traces are available from the corresponding author upon reasonable request. The custom Python-based wire myography analysis toolkit is publicly available at https://doi.org/10.5281/zenodo.18472624. No large-scale datasets requiring deposition in a public repository were generated in this study.

## SUPPLEMENTAL MATERIAL

Supplementary Tables S1-S5 are available online at the end of this document.

**Table S1**: Sex-specific analysis of Yoda2 effects on contractile parameters

**Table S2**: Effects of SR Ca²⁺ store depletion on Yoda2-mediated contractile inhibition

**Table S3**: Patch clamp electrophysiology data with pharmacological interventions

**Table S4**: Colocalization analysis of Piezo1 with RyR and BK channels

**Table S5**: Antibodies, reagents, chemicals and software source data

## ACKNOWLEDGMENTS

We would like to thank Brandon Light, Michele Persiani, and Camryn Sellers-Porter of the McElroy Lab for their technical support. We would also like to thank Rosie Dixon and Madeline Nieves-Cintron for their helpful discussions and suggestions.

## DISCLAIMERS

The content is solely the responsibility of the authors and does not necessarily represent the official views of the University of California, Davis.

## Supplementary Tables

**Table S1.**
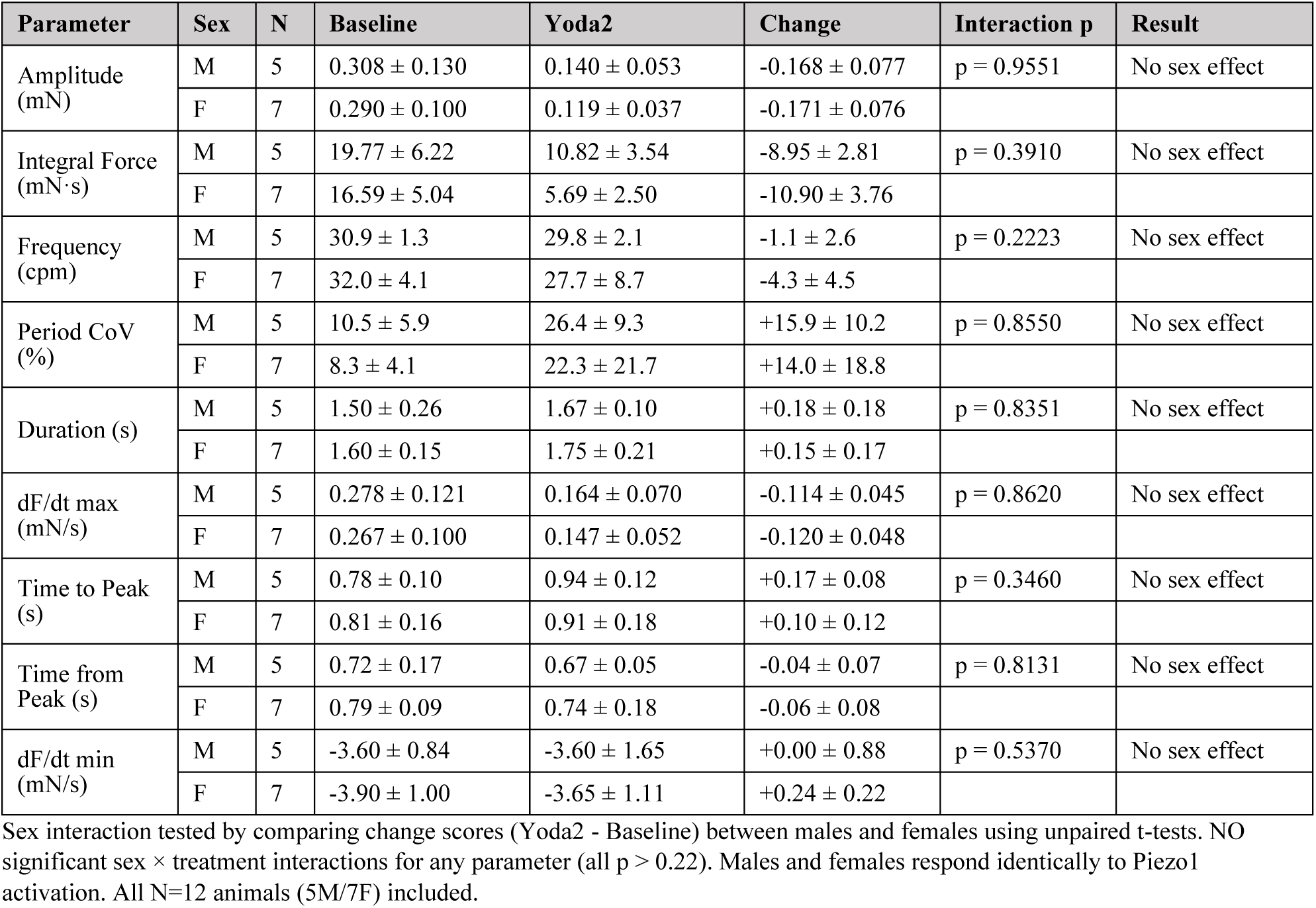
Sex-Disaggregated Analysis of Contractility Data.

**Table S2.**
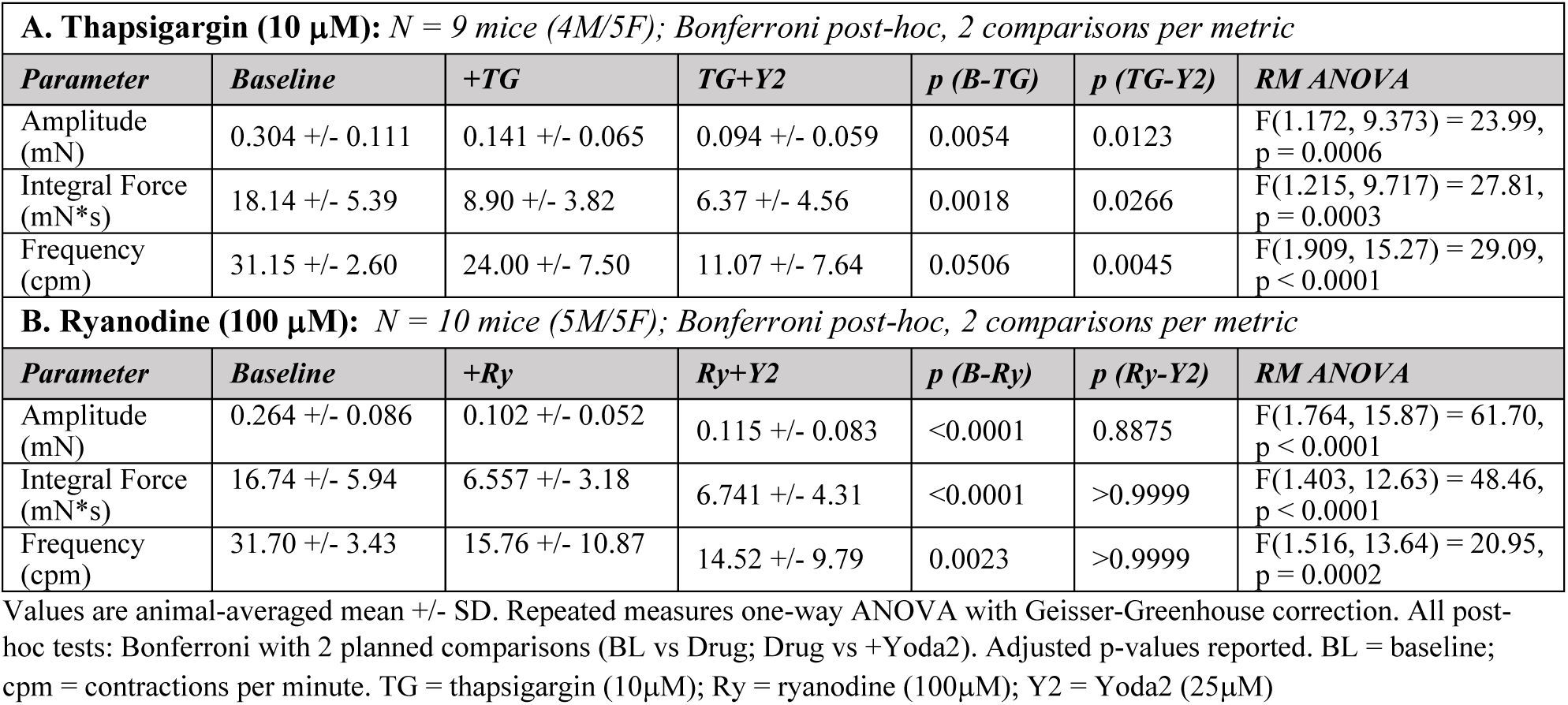
Contractile parameters during pharmacological SR disruption.

**Table S3.**
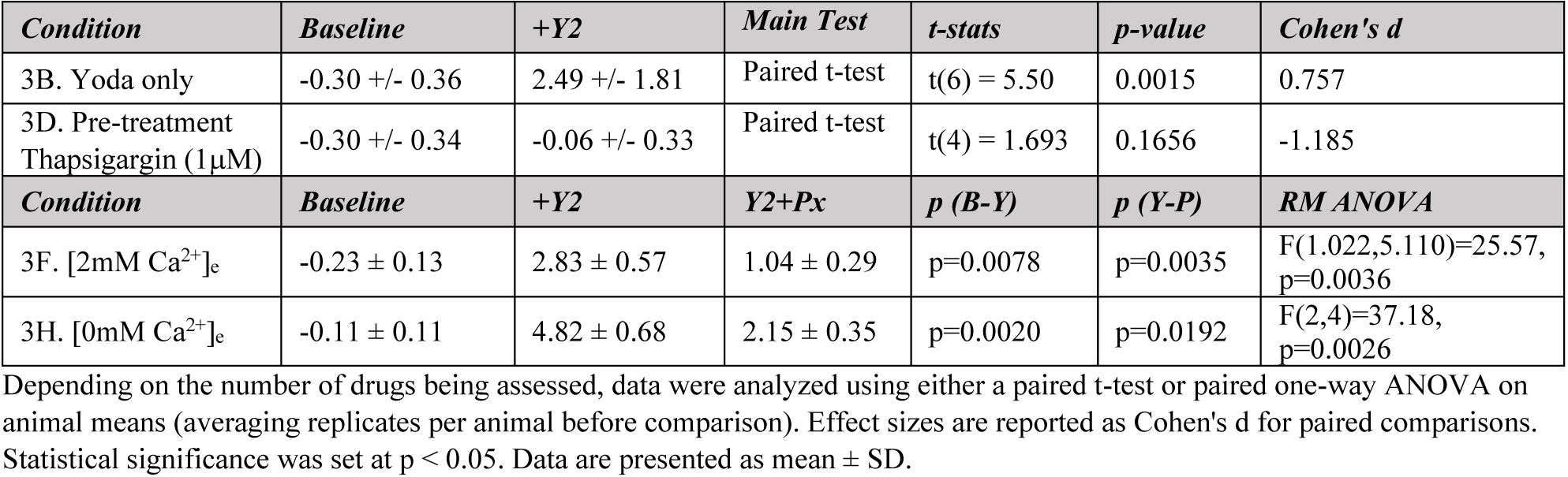
Whole Cell Patch Clamping Statistical Analysis.

**Table S4.**
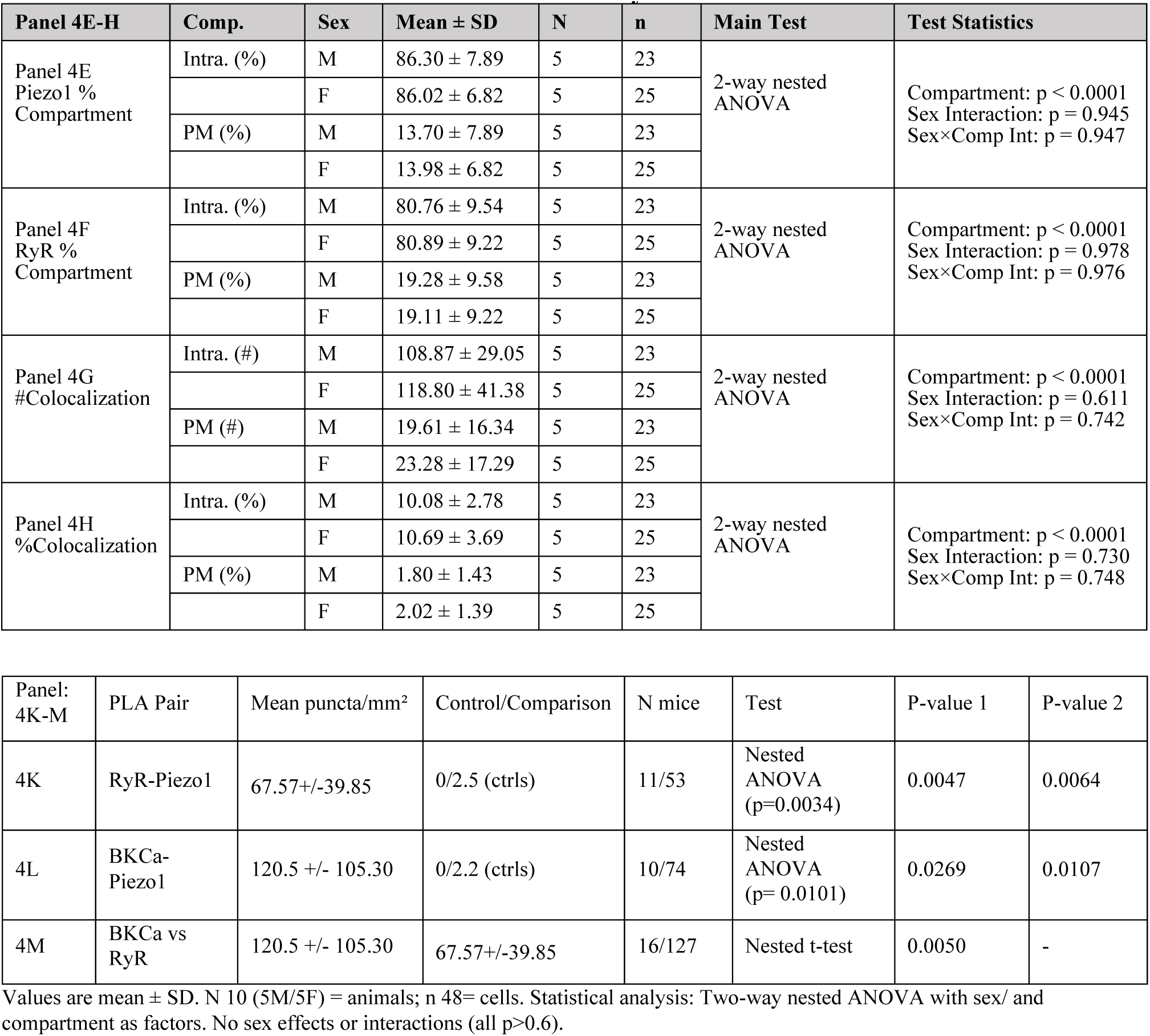
Subcellular Localization of Piezo1 and RyR and PLA Statistics.

**Table S5:**
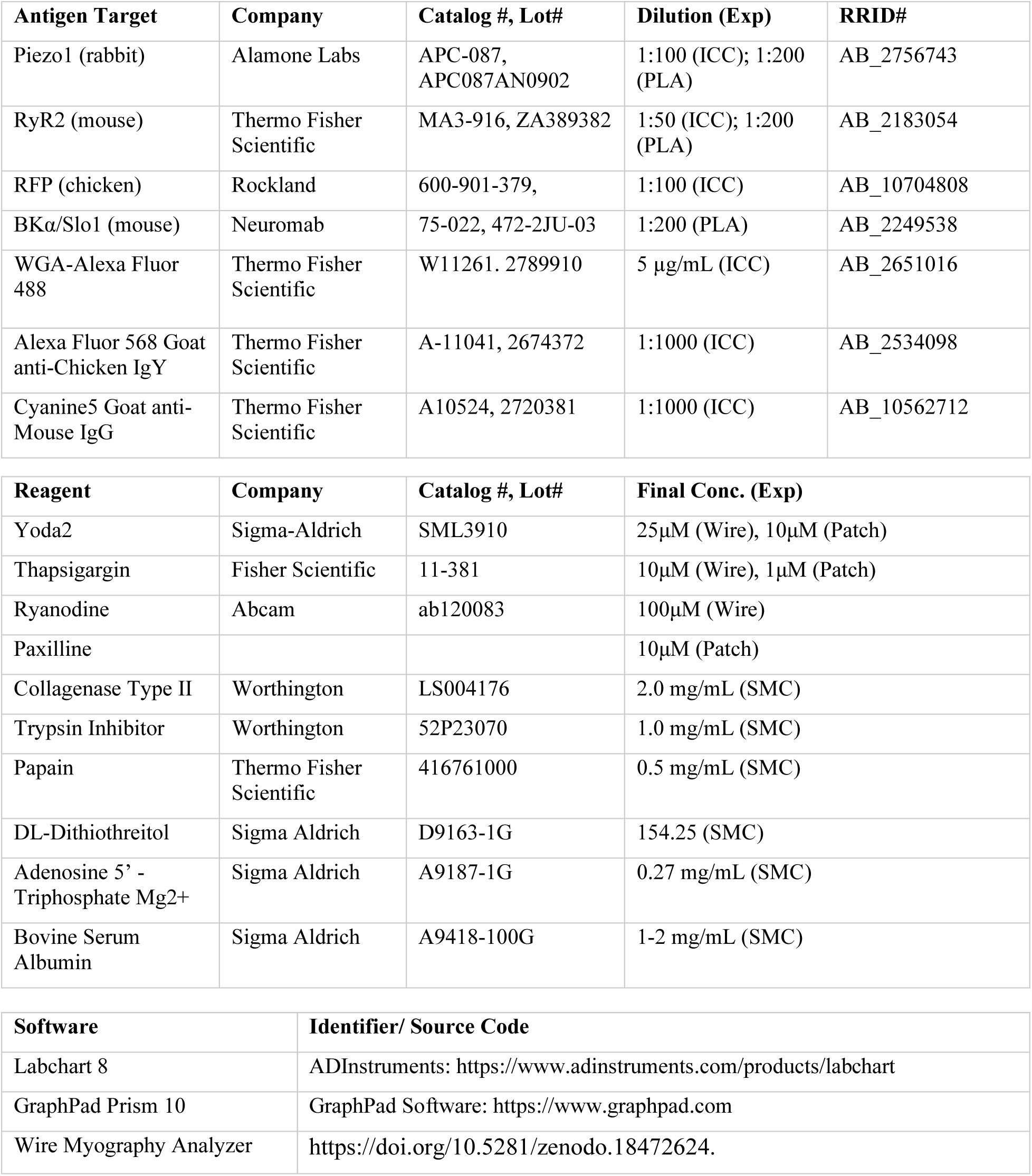
Antibodies, Reagents, Chemicals and Software Source Data.

| Reagent | Company | Catalog #, Lot# | Final Conc. (Exp) |
| --- | --- | --- | --- |
| Yoda2 | Sigma-Aldrich | SML3910 | 25μM (Wire), 10μM (Patch) |
| Thapsigargin | Fisher Scientific | 11-381 | 10μM (Wire), 1μM (Patch) |
| Ryanodine | Abcam | ab120083 | 100μM (Wire) |
| Paxilline | | | 10μM (Patch) |
| Collagenase Type II | Worthington | LS004176 | 2.0 mg/mL (SMC) |
| Trypsin Inhibitor | Worthington | 52P23070 | 1.0 mg/mL (SMC) |
| Papain | Thermo Fisher Scientific | 416761000 | 0.5 mg/mL (SMC) |
| DL-Dithiothreitol | Sigma Aldrich | D9163-1G | 154.25 (SMC) |
| Adenosine 5' - Triphosphate Mg<sup>2+</sup> | Sigma Aldrich | A9187-1G | 0.27 mg/mL (SMC) |
| Bovine Serum Albumin | Sigma Aldrich | A9418-100G | 1-2 mg/mL (SMC) |

