## Supplemental Tables S1-S5 for "Intracellular Mechanosensation in Intestinal Smooth Muscle: Piezo1 Complexes Amplify Signaling Beyond the Surface"

**Supplementary Tables**

| **Table S1. Sex-Disaggregated Analysis of Contractility Data** | | | | | | | |
| --- | --- | --- | --- | --- | --- | --- | --- |
| **Parameter** | **Sex** | **N** | **Baseline** | **Yoda2** | **Change** | **Interaction p** | **Result** |
| Amplitude (mN) | M | 5 | 0.308 ± 0.130 | 0.140 ± 0.053 | -0.168 ± 0.077 | p = 0.9551 | No sex effect |
|  | F | 7 | 0.290 ± 0.100 | 0.119 ± 0.037 | -0.171 ± 0.076 |  |  |
| Integral Force (mN·s) | M | 5 | 19.77 ± 6.22 | 10.82 ± 3.54 | -8.95 ± 2.81 | p = 0.3910 | No sex effect |
|  | F | 7 | 16.59 ± 5.04 | 5.69 ± 2.50 | -10.90 ± 3.76 |  |  |
| Frequency (cpm) | M | 5 | 30.9 ± 1.3 | 29.8 ± 2.1 | -1.1 ± 2.6 | p = 0.2223 | No sex effect |
|  | F | 7 | 32.0 ± 4.1 | 27.7 ± 8.7 | -4.3 ± 4.5 |  |  |
| Period CoV (%) | M | 5 | 10.5 ± 5.9 | 26.4 ± 9.3 | +15.9 ± 10.2 | p = 0.8550 | No sex effect |
|  | F | 7 | 8.3 ± 4.1 | 22.3 ± 21.7 | +14.0 ± 18.8 |  |  |
| Duration (s) | M | 5 | 1.50 ± 0.26 | 1.67 ± 0.10 | +0.18 ± 0.18 | p = 0.8351 | No sex effect |
|  | F | 7 | 1.60 ± 0.15 | 1.75 ± 0.21 | +0.15 ± 0.17 |  |  |
| dF/dt max (mN/s) | M | 5 | 0.278 ± 0.121 | 0.164 ± 0.070 | -0.114 ± 0.045 | p = 0.8620 | No sex effect |
|  | F | 7 | 0.267 ± 0.100 | 0.147 ± 0.052 | -0.120 ± 0.048 |  |  |
| Time to Peak (s) | M | 5 | 0.78 ± 0.10 | 0.94 ± 0.12 | +0.17 ± 0.08 | p = 0.3460 | No sex effect |
|  | F | 7 | 0.81 ± 0.16 | 0.91 ± 0.18 | +0.10 ± 0.12 |  |  |
| Time from Peak (s) | M | 5 | 0.72 ± 0.17 | 0.67 ± 0.05 | -0.04 ± 0.07 | p = 0.8131 | No sex effect |
|  | F | 7 | 0.79 ± 0.09 | 0.74 ± 0.18 | -0.06 ± 0.08 |  |  |
| dF/dt min (mN/s) | M | 5 | -3.60 ± 0.84 | -3.60 ± 1.65 | +0.00 ± 0.88 | p = 0.5370 | No sex effect |
|  | F | 7 | -3.90 ± 1.00 | -3.65 ± 1.11 | +0.24 ± 0.22 |  |  |

Sex interaction tested by comparing change scores (Yoda2 - Baseline) between males and females using unpaired t-tests. NO significant sex × treatment interactions for any parameter (all p > 0.22). Males and females respond identically to Piezo1 activation. All N=12 animals (5M/7F) included.

| **Table S2. Contractile parameters during pharmacological SR disruption.** | | | | | | |
| --- | --- | --- | --- | --- | --- | --- |
| **A. Thapsigargin (10 μM):** *N = 9 mice (4M/5F); Bonferroni post-hoc, 2 comparisons per metric* | | | | | | |
| ***Parameter*** | ***Baseline*** | ***+TG*** | ***TG+Y2*** | ***p (B-TG)*** | ***p (TG-Y2)*** | ***RM ANOVA*** |
| Amplitude (mN) | 0.304 +/- 0.111 | 0.141 +/- 0.065 | 0.094 +/- 0.059 | 0.0054 | 0.0123 | F(1.172, 9.373) = 23.99,  p = 0.0006 |
| Integral Force (mN*s) | 18.14 +/- 5.39 | 8.90 +/- 3.82 | 6.37 +/- 4.56 | 0.0018 | 0.0266 | F(1.215, 9.717) = 27.81,  p = 0.0003 |
| Frequency (cpm) | 31.15 +/- 2.60 | 24.00 +/- 7.50 | 11.07 +/- 7.64 | 0.0506 | 0.0045 | F(1.909, 15.27) = 29.09,  p < 0.0001 |
| **B. Ryanodine (100 μM):** *N = 10 mice (5M/5F); Bonferroni post-hoc, 2 comparisons per metric* | | | | | | |
| ***Parameter*** | ***Baseline*** | ***+Ry*** | ***Ry+Y2*** | ***p (B-Ry)*** | ***p (Ry-Y2)*** | ***RM ANOVA*** |
| Amplitude (mN) | 0.264 +/- 0.086 | 0.102 +/- 0.052 | 0.115 +/- 0.083 | <0.0001 | 0.8875 | F(1.764, 15.87) = 61.70, p < 0.0001 |
| Integral Force (mN*s) | 16.74 +/- 5.94 | 6.557 +/- 3.18 | 6.741 +/- 4.31 | <0.0001 | >0.9999 | F(1.403, 12.63) = 48.46, p < 0.0001 |
| Frequency (cpm) | 31.70 +/- 3.43 | 15.76 +/- 10.87 | 14.52 +/- 9.79 | 0.0023 | >0.9999 | F(1.516, 13.64) = 20.95, p = 0.0002 |

Values are animal-averaged mean +/- SD. Repeated measures one-way ANOVA with Geisser-Greenhouse correction. All post-hoc tests: Bonferroni with 2 planned comparisons (BL vs Drug; Drug vs +Yoda2). Adjusted p-values reported. BL = baseline; cpm = contractions per minute. TG = thapsigargin (10μM); Ry = ryanodine (100μM); Y2 = Yoda2 (25μM)

| **Table S3. Whole Cell Patch Clamping Statistical Analysis** | | | | | | |
| --- | --- | --- | --- | --- | --- | --- |
| ***Condition*** | ***Baseline*** | ***+Y2*** | ***Main Test*** | ***t-stats*** | ***p-value*** | ***Cohen's d*** |
| 3B. Yoda only | -0.30 +/- 0.36 | 2.49 +/- 1.81 | Paired t-test | t(6) = 5.50 | 0.0015 | 0.757 |
| 3D. Pre-treatment Thapsigargin (1μM) | -0.30 +/- 0.34 | -0.06 +/- 0.33 | Paired t-test | t(4) = 1.693 | 0.1656 | -1.185 |
| ***Condition*** | ***Baseline*** | ***+Y2*** | ***Y2+Px*** | ***p (B-Y)*** | ***p (Y-P)*** | ***RM ANOVA*** |
| 3F. [2mM Ca^2+^]_e_ | -0.23 ± 0.13 | 2.83 ± 0.57 | 1.04 ± 0.29 | p=0.0078 | p=0.0035 | F(1.022,5.110)=25.57, p=0.0036 |
| 3H. [0mM Ca^2+^]_e_ | -0.11 ± 0.11 | 4.82 ± 0.68 | 2.15 ± 0.35 | p=0.0020 | p=0.0192 | F(2,4)=37.18, p=0.0026 |

Depending on the number of drugs being assessed, data were analyzed using either a paired t-test or paired one-way ANOVA on animal means (averaging replicates per animal before comparison). Effect sizes are reported as Cohen's d for paired comparisons. Statistical significance was set at p < 0.05. Data are presented as mean ± SD.

| **Table S4. Subcellular Localization of Piezo1 and RyR and PLA Statistics** | | | | | | | |
| --- | --- | --- | --- | --- | --- | --- | --- |
| **Panel 4E-H** | **Comp.** | **Sex** | **Mean ± SD** | **N** | **n** | **Main Test** | **Test Statistics** |
| Panel 4E  Piezo1 %  Compartment | Intra. (%) | M | 86.30 ± 7.89 | 5 | 23 | 2-way nested  ANOVA | Compartment: p < 0.0001  Sex Interaction: p = 0.945  Sex×Comp Int: p = 0.947 |
|  |  | F | 86.02 ± 6.82 | 5 | 25 |  |  |
|  | PM (%) | M | 13.70 ± 7.89 | 5 | 23 |  |  |
|  |  | F | 13.98 ± 6.82 | 5 | 25 |  |  |
| Panel 4F  RyR %  Compartment | Intra. (%) | M | 80.76 ± 9.54 | 5 | 23 | 2-way nested  ANOVA | Compartment: p < 0.0001  Sex Interaction: p = 0.978  Sex×Comp Int: p = 0.976 |
|  |  | F | 80.89 ± 9.22 | 5 | 25 |  |  |
|  | PM (%) | M | 19.28 ± 9.58 | 5 | 23 |  |  |
|  |  | F | 19.11 ± 9.22 | 5 | 25 |  |  |
| Panel 4G  #Colocalization | Intra. (#) | M | 108.87 ± 29.05 | 5 | 23 | 2-way nested  ANOVA | Compartment: p < 0.0001  Sex Interaction: p = 0.611  Sex×Comp Int: p = 0.742 |
|  |  | F | 118.80 ± 41.38 | 5 | 25 |  |  |
|  | PM (#) | M | 19.61 ± 16.34 | 5 | 23 |  |  |
|  |  | F | 23.28 ± 17.29 | 5 | 25 |  |  |
| Panel 4H  %Colocalization | Intra. (%) | M | 10.08 ± 2.78 | 5 | 23 | 2-way nested  ANOVA | Compartment: p < 0.0001  Sex Interaction: p = 0.730  Sex×Comp Int: p = 0.748 |
|  |  | F | 10.69 ± 3.69 | 5 | 25 |  |  |
|  | PM (%) | M | 1.80 ± 1.43 | 5 | 23 |  |  |
|  |  | F | 2.02 ± 1.39 | 5 | 25 |  |  |

| Panel: 4K-M | PLA Pair | Mean puncta/mm² | Control/Comparison | N mice | Test | P-value 1 | P-value 2 |
| --- | --- | --- | --- | --- | --- | --- | --- |
| 4K | RyR-Piezo1 | 67.57+/-39.85 | 0/2.5 (ctrls) | 11/53 | Nested ANOVA (p=0.0034) | 0.0047 | 0.0064 |
| 4L | BKCa-Piezo1 | 120.5 +/- 105.30 | 0/2.2 (ctrls) | 10/74 | Nested ANOVA  (p= 0.0101) | 0.0269 | 0.0107 |
| 4M | BKCa vs RyR | 120.5 +/- 105.30 | 67.57+/-39.85 | 16/127 | Nested t-test | 0.0050 | - |

Values are mean ± SD. N 10 (5M/5F) = animals; n 48= cells. Statistical analysis: Two-way nested ANOVA with sex/ and compartment as factors. No sex effects or interactions (all p>0.6).

**Table S5: Antibodies, Reagents, Chemicals and Software Source Data**

| **Antigen Target** | **Company** | **Catalog #, Lot#** | **Dilution (Exp)** | **RRID#** |
| --- | --- | --- | --- | --- |
| Piezo1 (rabbit) | Alamone Labs | APC-087, APC087AN0902 | 1:100 (ICC); 1:200 (PLA) | AB_2756743 |
| RyR2 (mouse) | Thermo Fisher Scientific | MA3-916, ZA389382 | 1:50 (ICC); 1:200 (PLA) | AB_2183054 |
| RFP (chicken) | Rockland | 600-901-379, | 1:100 (ICC) | AB_10704808 |
| BKα/Slo1 (mouse) | Neuromab | 75-022, 472-2JU-03 | 1:200 (PLA) | AB_2249538 |
| WGA-Alexa Fluor 488 | Thermo Fisher Scientific | W11261. 2789910 | 5 µg/mL (ICC) | AB_2651016 |
| Alexa Fluor 568 Goat anti-Chicken IgY | Thermo Fisher Scientific | A-11041, 2674372 | 1:1000 (ICC) | AB_2534098 |
| Cyanine5 Goat anti-Mouse IgG | Thermo Fisher Scientific | A10524, 2720381 | 1:1000 (ICC) | AB_10562712 |

| **Reagent** | **Company** | **Catalog #, Lot#** | **Final Conc. (Exp)** |
| --- | --- | --- | --- |
| Yoda2 | Sigma-Aldrich | SML3910 | 25μM (Wire), 10μM (Patch) |
| Thapsigargin | Fisher Scientific | 11-381 | 10μM (Wire), 1μM (Patch) |
| Ryanodine | Abcam | ab120083 | 100μM (Wire) |
| Paxilline |  |  | 10μM (Patch) |
| Collagenase Type II | Worthington | LS004176 | 2.0 mg/mL (SMC) |
| Trypsin Inhibitor | Worthington | 52P23070 | 1.0 mg/mL (SMC) |
| Papain | Thermo Fisher Scientific | 416761000 | 0.5 mg/mL (SMC) |
| DL-Dithiothreitol | Sigma Aldrich | D9163-1G | 154.25 (SMC) |
| Adenosine 5’ -Triphosphate Mg2+ | Sigma Aldrich | A9187-1G | 0.27 mg/mL (SMC) |
| Bovine Serum Albumin | Sigma Aldrich | A9418-100G | 1-2 mg/mL (SMC) |

| **Software** | **Identifier/ Source Code** |
| --- | --- |
| Labchart 8 | ADInstruments: https://www.adinstruments.com/products/labchart |
| GraphPad Prism 10 | GraphPad Software: https://www.graphpad.com |
| Wire Myography Analyzer | https://doi.org/10.5281/zenodo.18472624. |
